# Geometric changes in the nucleoids of Deinococcus radiodurans reveal involvement of new proteins in recovery from ionizing radiation

**DOI:** 10.1101/2024.01.17.576117

**Authors:** Antonio Cordova, Brandon Niese, Philip Sweet, Pratik Kamat, Jude M Phillip, Vernita Gordon, Lydia M. Contreras

## Abstract

The extremophile *Deinococcus radiodurans* maintains a highly-organized and condensed nucleoid as its default state, possibly contributing to high tolerance of ionizing radiation (IR). Previous studies of the *D. radiodurans* nucleoid were limited by reliance on manual image annotation and qualitative metrics. Here, we introduce a high-throughput approach to quantify the geometric properties of cells and nucleoids, using confocal microscopy, digital reconstructions of cells, and computational modeling. We utilize this novel approach to investigate the dynamic process of nucleoid condensation in response to IR stress. Our quantitative analysis reveals that at the population level, exposure to IR induced nucleoid compaction and decreased size of *D. radiodurans* cells. Morphological analysis and clustering identified six distinct sub-populations across all tested experimental conditions. Results indicate that exposure to IR induces fractional redistributions of cells across sub-populations to exhibit morphologies that associate with greater nucleoid condensation, and decreased abundance of sub-populations associated with cell division. Nucleoid associated proteins (NAPs) may link nucleoid compaction and stress tolerance, but their roles in regulating compaction in *D. radiodurans* is unknown. Imaging of genomic mutants of known and suspected NAPs that contribute to nucleoid condensation found that deletion of nucleic acid binding proteins, not previously described as NAPs, can remodel the nucleoid by driving condensation or decondensation in the absence of stress and that IR increases the abundance of these morphological states. Thus, our integrated analysis introduces a new methodology for studying environmental influences on bacterial nucleoids and provides an opportunity to further investigate potential regulators of nucleoid condensation.

**Importance:** *D. radiodurans*, an extremophile known for its stress tolerance, constitutively maintains a highly-condensed nucleoid. Qualitative studies have described nucleoid behavior under a variety of conditions. However, a lack of quantitative data regarding nucleoid organization and dynamics have limited our understanding of regulatory mechanisms controlling nucleoid organization in *D. radiodurans*. Here, we introduce a quantitative approach that enables high-throughput quantitative measurements of subcellular spatial characteristics in bacterial cells. Applying this to wild-type or single-protein-deficient populations of *D. radiodurans* subjected to ionizing radiation, we identified significant stress-responsive changes in cell shape, nucleoid organization, and morphology. These findings highlight this methodology’s adaptability and capacity for quantitatively analyzing the cellular response to stressors for screening cellular proteins involved in bacterial nucleoid organization.

## Introduction

The compaction of the chromosome into a hierarchically organized irregularly-shaped nucleoid is one example of subcellular spatial organization in bacteria [1, 2]. Nucleoids contain all or most of the genetic materials of the bacterial cell and are essential to gene expression and cell division [1]. Regulation of the structural organization and the morphologies of bacterial nucleoids, are best understood only in a few model organisms, such as *Escherichia coli* and *Bacillus subtilis* [3]. Furthermore, molecular mechanisms that explain the dynamics of how nucleoids change shapes under different cellular and environmental conditions are not well understood, particularly in the context of stress-response regulation.

Since genomic material requires compaction to fit into the cell, it has been proposed that multiple classes of cellular factors and mechanisms are involved in forming the bacterial nucleoid. These classes include nucleoid-binding proteins, non-coding nucleoid-associated RNAs [4], and DNA supercoiling mechanisms [5]. Nucleoid-associated proteins (NAP) are suspected to contribute to DNA-DNA contacts that promote higher-order organization. For example, the *E.*c*oli* NAPs histone-like nucleoid structuring protein (H-NS), ParB, and structural maintenance of chromosomes (SMC) facilitate DNA bridging and higher-order organization [6]. Likewise, histone-like protein (HU) binds to and bends DNA to maintain the nucleoid [7]. In addition to protein factors, a study of *E.coli* has identified the presence of a noncoding RNA (naRNA4) in the nucleoid fraction; it has been suggested that naRNA4 may, in associations with nucleoid-associated proteins (like HU), contribute to DNA compaction by forming more remote DNA-DNA connections [8].

In addition to providing structure to the nucleoid, NAPs can contribute to stress tolerance. For examples, HU homologs in extremophilic bacteria are suspected to play a role in survival under radiation-based stressors [9] and the *E. coli* Dps protein, which protects DNA from oxidative damage, also contributes to DNA condensation [10, 11].

Earlier *E.coli* studies have shown that the nucleoid structure is dynamic and responsive to both changes in the physiological state of the cell [2] and to changes in environmental conditions [12]. For instance, oxidative stress that threatens DNA integrity has been shown to induce nucleoid compaction [13]. In *B. subtilis*, a qualitative study found that under non-stressed conditions the nucleoid occupied most of the cell volume, and that non-lethal exposures to UV light resulted in compacted chromosomes and elongated cells [14]. This change in nucleoid morphology is reversible, particularly after lower doses of UV, in a way that depends on active mechanisms of DNA repair. These findings suggest that this case of nucleoid compaction might be correlated with active and controlled DNA repair and/or to mechanisms of DNA protection under stress. For instance, in *E. coli*, RecA is a key protein associated with DNA repair, and the association of DNA with the active form of RecA affects nucleoid compaction [12]. In similar studies, damage to DNA and/or disruptions to DNA repair induced in *E.coli*, *Mycobacterium smegmatis* [15] and *Staphlyococcus aureus* [13] also led to compaction of the genome. Specifically, studies on DNA damage stress in *E. coli* found the bacterial chromatin compacted down to fill ∼20% of the cell volume when exposed to nalidixic acid, a double-strand DNA damage agent, as opposed to occupying the majority of the cytoplasmic, ribosome-free space, under no stress [12]. In summary, many studies across many organisms have suggested that the volume fraction of the cell occupied by the nucleoid and how the nucleoid reconfigures after stress are both of biological relevance.

In most bacteria, including *E. coli* and *B. subtilis,* the D_10_ value (the dose of ionizing radiation (IR) required to reduce a bacterial population to 10% of its initial size) is less than 1 kGy [16]. In stark contrast, *D. radiodurans* has high radiation tolerance (D_10_ value of ∼12.5 kGy) [17]. Surprisingly, little is known about the types of unique mechanisms that *D. radiodurans* uses in protecting and coordinating repair of its genome under stress [18–21].

Characteristics that have been suggested as possibly contributing to *D. radiodurans*’ tolerance of oxidative stressors, such as IR, include its robust DNA repair machinery, the Signal Recognition Particle pathway of antioxidant protein transport, its IR-sensitive post-transcriptional regulator network, and its constitutively compacted genome [22–28]. The last is of particular importance for the work we present here. Examining the nucleoid structure and structural dynamics in an organism, like *D. radiodurans*, that maintains a highly-organized, condensed nucleoid by default, has the potential to shed light on how subcellular organization is regulated [29]. Understanding how the organization and response dynamics of the condensed *D. radiodurans* nucleoid contributes to its high IR tolerance, and how this depends on the activity of DNA-binding proteins and nucleoid-associated proteins, has the potential to provide a unique perspective on subcellular organization in prokaryotes.

Qualitative studies of *D. radiodurans* [29–32] have confirmed that nucleoid morphology is highly dynamic during the cell cycle, even in the absence of any environmental stresses. Prior qualitative descriptions of the *D. radiodurans* nucleoid under stress have been based on fluorescent microscopy and on cryo-electron microscopy [31, 32]. These earlier, qualitative studies point to the need for deeper investigation of the dynamics of nucleoids. Deeper investigation requires methods for quantitative image analysis with the ability to measure many types of geometric properties that can be defined by investigators *ad hoc* to better map the dynamics of nucleoids. Ideally, such an approach would also be high-throughput, able to quickly analyze large numbers of cells as well as many different geometric parameters.

Here, we develop a method for quantitative measurement of geometric properties of bacterial cells and nucleoids. We applied this method to populations of *Deinococcus radiodurans* to study the impact of IR and the contributions of different nucleic acid binding proteins on nucleoid structure. Our approach uses machine learning to create analyzable digital re-creations of individual bacterial cells from laser-scanning confocal microscopy images of cells pre and post IR exposure. Digital re-creations are input into Python code for analysis, so that any user-defined parameter can be measured. Following either IR exposure or the deletion of single proteins, we measure statistically-significant changes in spatial and geometric parameters of *D. radiodurans*. Consistent with previous observations of bacteria under DNA-damaging stress, we find that exposure to IR results in decreased cell area while also resulting in more circular, less eccentric nucleoids. Our quantitative method also allows us to measure previously-unreported shifts in the nucleoid area and the fraction of the cell occupied by the nucleoid. We track these dose-dependent geometric parameters over 18 hours to measure the dynamics of recovery after exposure to IR and we show that the recovery process also varies with initial, acute dose.

In tandem with our quantitative method, our morphological clustering analysis identifies extremes in cell and nucleoid morphology that can effectively assist in determining bacterial sub-populations across experimental conditions. We find that exposure to IR decreases the representation of sub-populations with morphological features associated with cell division. Additionally, we find that IR stress increases the representation of sub-populations with morphologies associated with greater nucleoid condensation. We find that the representation of these morphological clusters can be driven by deletion of specific proteins in *D. radiodurans,* which we hypothesize contribute to mechanisms of nucleoid dynamics under stress.

Specifically, we identified DNA-binding, RNA-binding, and stress-response proteins that were not previously identified as nucleoid-associated to have significant effects on nucleoid remodeling under sham (no radiation) conditions. We also observe that sub-populations, representative of morphological states of nucleoid condensation or decondensation in the absence of stress, were significantly increased when cells were exposed to IR. Collectively, our results indicate that this approach can be used to screen candidate proteins for their role in IR-tolerance and -recovery. Overall, this integrated analysis introduces a novel approach for measuring changes in cell and nucleoid geometry, and for the investigation of key genes, known or unknown, in nucleoid condensation.

## Results

### Confocal laser-scanning microscopy enables qualitative capture of the characteristics of D. radiodurans nucleoids

To establish quantitative measures for the geometric properties of *D. radiodurans* cells and nucleoids, we first characterized a wild-type strain (R1) in the presence and the absence of ionizing radiation. The experimental scheme we used is illustrated in Figure 1A and described in more detail in the Methods section. In brief, bacterial cultures were exposed to different levels of ionizing radiation (0-12 kGy); 12 kGy was selected because *D. radiodurans* has a D10 value of ∼12.5 kGy [19]. Following irradiation and a 2-hour recovery period at 30C, cells were stained with two fluorescent dyes, Nile Red and Syto 9 Green, which associate with the hydrophobic region of the bacterial cell membrane and the bacterial DNA respectively. The two dyes Nile Red and Syto 9 Green have two different spectra for fluorescence excitation and emission, allowing the cell membrane (red) and the nucleoid (green) to be differentiated by color. We used laser-scanning confocal microscopy to image stained bacteria in these two separate image channels (Figure 1B).

**Figure 1.**
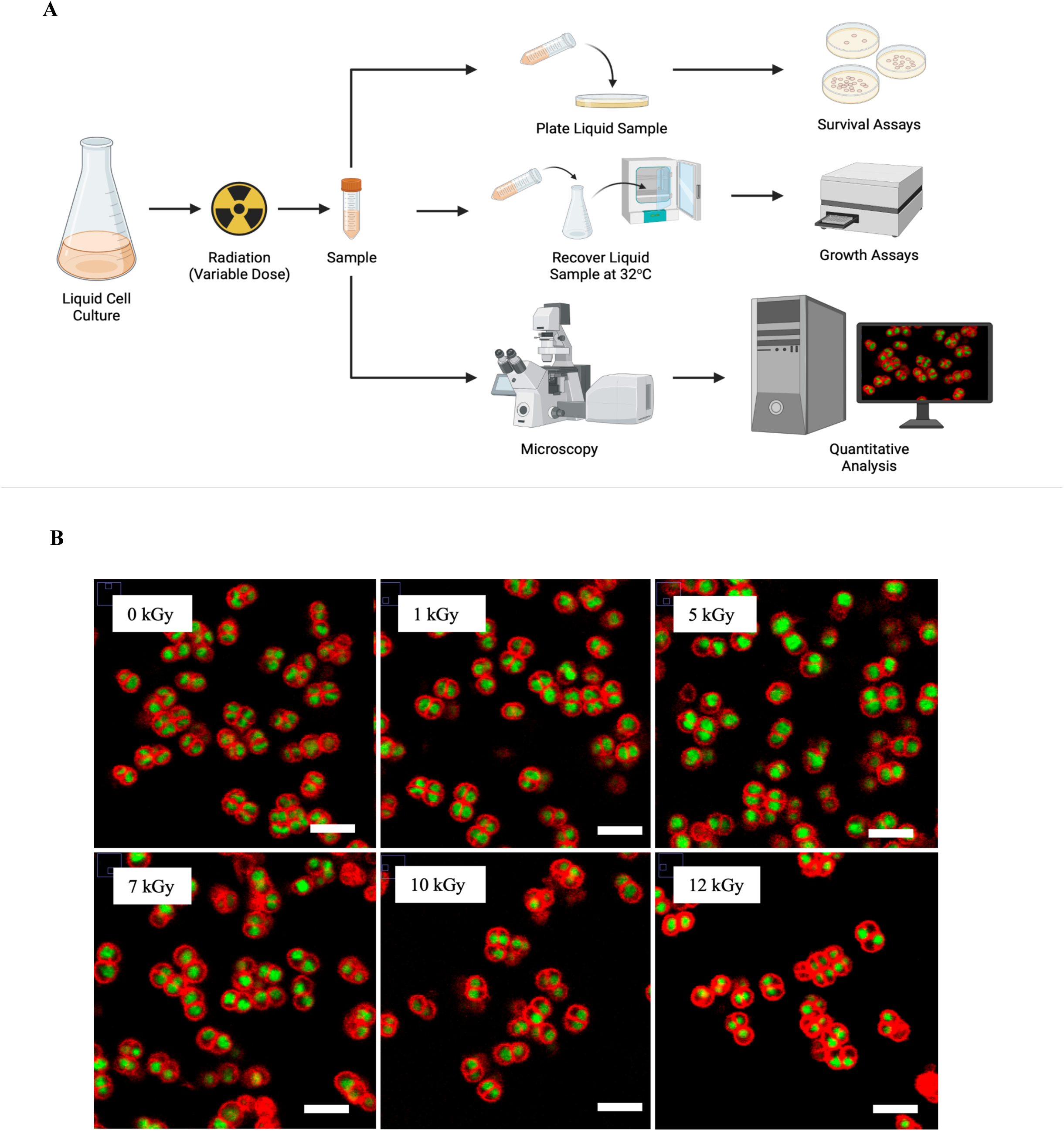
Experiment and analysis scheme, with confocal micrographs to illustrate the raw material for analysis. **(A)** Experimental scheme. *D. radiodurans* cultures were exposed to IR and then imaged using confocal microscopy. Created in BioRender. **(B)** Confocal micrographs of wild-type (R1) *D. radiodurans*, with cell membranes stained red and nucleoids stained green. Before imaging, the cells shown were exposed to 0, 1, 5, 7, 10, and 12 kGy radiation, as indicated on sub-panels. Scale bars are 5 μm.

Characterization of *D. radiodurans* survival and recovery after irradiation was performed by counting colony-forming units (CFU) to measure survival and by quantifying optical density over time to measure growth (Supplementary Figure S1 A-C). Our data confirms a decrease in viable surviving bacteria and a decrease in growth rate with IR exposure doses approaching 10 kGy or greater, as observed in our previous work [26].

In *D. radiodurans* cells that were not exposed to radiation, nucleoids are condensed and irregularly-shaped (Figure 1B) and membranes have a roughly circular cross-section (with flattening where they adjoin other cells in *D. radiodurans*’ characteristic tetrad arrangement). These findings, *i.e.* constitutively-condensed nucleoids inside quasi-spherical cells that are arranged in tetrads, agree well with published observations about *D. radiodurans*’ baseline state in the absence of radiation or other stressors [32].

Importantly, during recovery post exposure to ionizing radiation, cells trend toward more-circular nucleoids with increasing levels of radiation (Figure 1B). These qualitative trends are consistent with previous reports of more condensed nucleoids in *D. radiodurans* following radiation stress [25].

### Development of a quantitative framework for reliably capturing shifts in geometric characteristics of cells

Building on these qualitative observations, we next created a set of quantitative metrics that describe the geometric properties of cells and nucleoids and the relationship between both properties. Specifically, we developed a pipeline for quantitative image analysis, beginning with laser-scanning confocal fluorescence images, as shown in Figure 1B. For every micrograph, both the membrane and the nucleoid grayscale image channels were segmented using the Cellpose [33] machine learning segmentation library. Segmented regions were used to create masks, with each mask showing the fluorescence “footprint” of either one cell membrane or one nucleoid. Using position information, each cell’s membrane and nucleoid masks were matched together, with the criterion that there is only one nucleoid in each cell. A schematic cartoon outlining the key steps of the image analysis pipeline is shown in Figure 2A.

**Figure 2.**
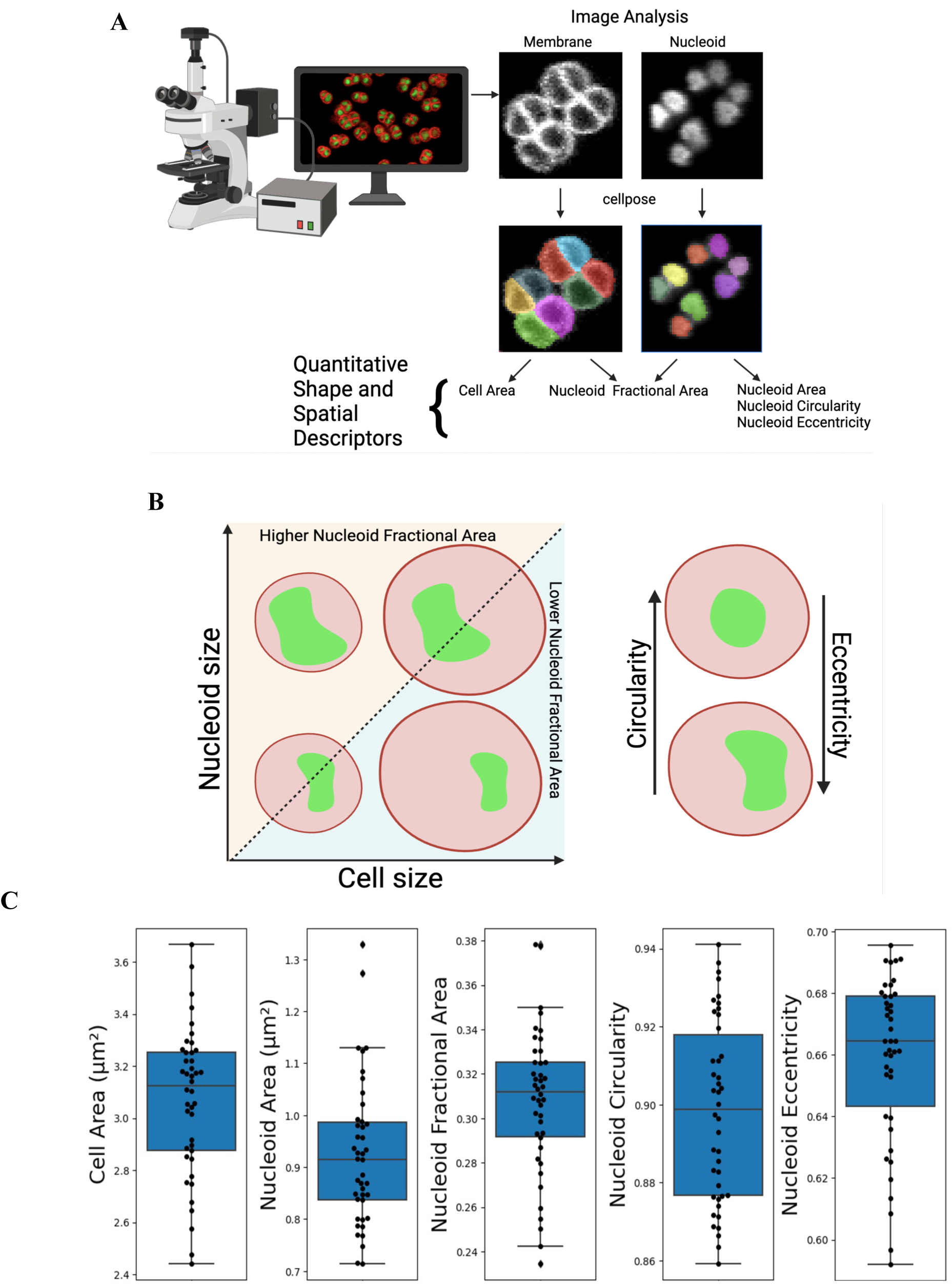
Image analysis for high-throughput quantification of geometric characteristics of cells and nucleoids. **(A)** The image analysis pipeline begins with laser-scanning confocal fluorescence micrographs, with cell membranes and nucleoids in different image channels. Micrographs are input into Cellpose, which creates segmented binary masks of each cell membrane and each nucleoid. In this figure, distinct binary masks are indicated with different colors. Computational analysis of masks is used to obtain user-defined geometric characteristics. Created in BioRender. **(B)** The parameter space examined in this study includes spatial and geometric characteristics. Spatial parameters include cell size, nucleoid size, and nucleoid fractional area. Geometric characteristics include nucleoid circularity and eccentricity. Created in BioRender. **(C)** Box-and-whisker plots of per-replicate mean values of geometric and spatial parameters for wild-type (R1) *D. radiodurans* cells that have not been irradiated. Means from 24 biological replicate populations are shown (black data points). Boxes show the middle 50% of the means, with horizontal lines representing the median value. Upper/lower whiskers represent upper/lower quartiles. An average of 1675 cells (min-max 262-3281) were measured for each biological replicate; 42 biological replicates. For each geometric characteristic measured, probability density distributions are shown in Supplementary Figures S2-S5.

#### Defining Bulk Cell Metric**s**

Five metrics were used to describe the cells: Cell Area, Nucleoid Area, Nucleoid Fractional Area, Nucleoid Circularity, and Nucleoid Eccentricity. The area enclosed by the membrane mask was used to measure the cross-sectional area of each cell (Cell Area), and the area covered by the nucleoid mask was used to measure the cross-sectional area of each nucleoid (Nucleoid Area). Measuring cell area allows assessment of spatial changes in cells that are not directly related to any change in nucleoid properties but are likely to arise from radiative damage [34]. Given that changes in relative cell and nucleoid area sizes are easily conflated with changes in absolute size following qualitative inspection by the human eye, we provide a quantitative measure of relative cell-nucleoid sizes by introducing the parameter nucleoid fractional area (NFA); this is the ratio of nucleoid cross-sectional area to cell cross-sectional area, using the matched pairs of each cell’s membrane and nucleoid masks. Nucleoid fractional area provides a quantitative measure of the nucleoid size relative to the cell size. A conceptual cartoon of the relationship between cell size, nucleoid size, and nucleoid fractional area is shown in Figure 2B (left side). In addition to differences in cell and nucleoid size, visual inspection of micrograph images (Figure 1B) shows differences in the cross-sectional shapes of nucleoids. We measure nucleoid shape using Circularity and Eccentricity (Figure 2B).

Using these geometric parameters (the calculations of which are detailed in the Methods section) we have started by characterizing non-irradiated *D. radiodurans* (Figure 2C and Supplementary Figures S2-S5). Specifically, we have found that, in the absence of radiation stress, wild-type *D. radiodurans* cells have a mean cross-sectional area of about 3 *μm* and their nucleoids have a mean cross-sectional area of about 0.9 *μm*. Their mean nucleoid fractional area is slightly less than 0.3. The mean nucleoid circularity is about 0.9 and the mean nucleoid eccentricity is about 0.7. Variations across biological replicas across different days are included as empirically-measured probability density distributions (Supplementary Figures S2-S5).

Kolmogorov-Smirnov testing (K-S test) was used to determine statistically-significant differences associated with irradiation in almost all cases (Supplementary Figure S2-S5, Supplementary Table S1), as it can compare distributions describing individuals in the entire measured population. K-S tests can exhibit sensitivity to changes in the distribution of a small number of outliers; however, this does not necessarily reflect overall changes in the population.

### Bulk quantitative measurements of changes in cellular characteristics during recovery post exposure to ionizing radiation indicate nucleoid compaction upon radiation exposure

We characterized cell and nucleoid geometries in the wild-type R1 strain, in the presence of radiation, and compared these to the geometries measured for the wild-type R1 strain in the absence of radiation. Observed changes in the geometric parameters are shown in Figure 3A (Supplementary Figure S6), and the results of significance testing are shown in Supplementary Table S3. We varied ionizing radiation levels from 0 to 12 kGy. A maximum acute dose of 12 kGy was used to approach the D10 value of ∼12.5 kGy for *D. radiodurans*. For the measurement of radiation-induced changes while accounting for inter-replicate variation, we record the arithmetic mean of each value, for the paired irradiated and non-irradiated biological replicates. For each day’s replicate, the ratio of the irradiated value to the non-irradiated value measures the fold change in that value. This was done for each replicate, for a total of six measurements of fold changes for each characteristic. This normalization approach is intended to account for day-to-day variations, and it parallels day-by-day normalization approaches we have used in previous work [35, 36].

**Figure 3.**
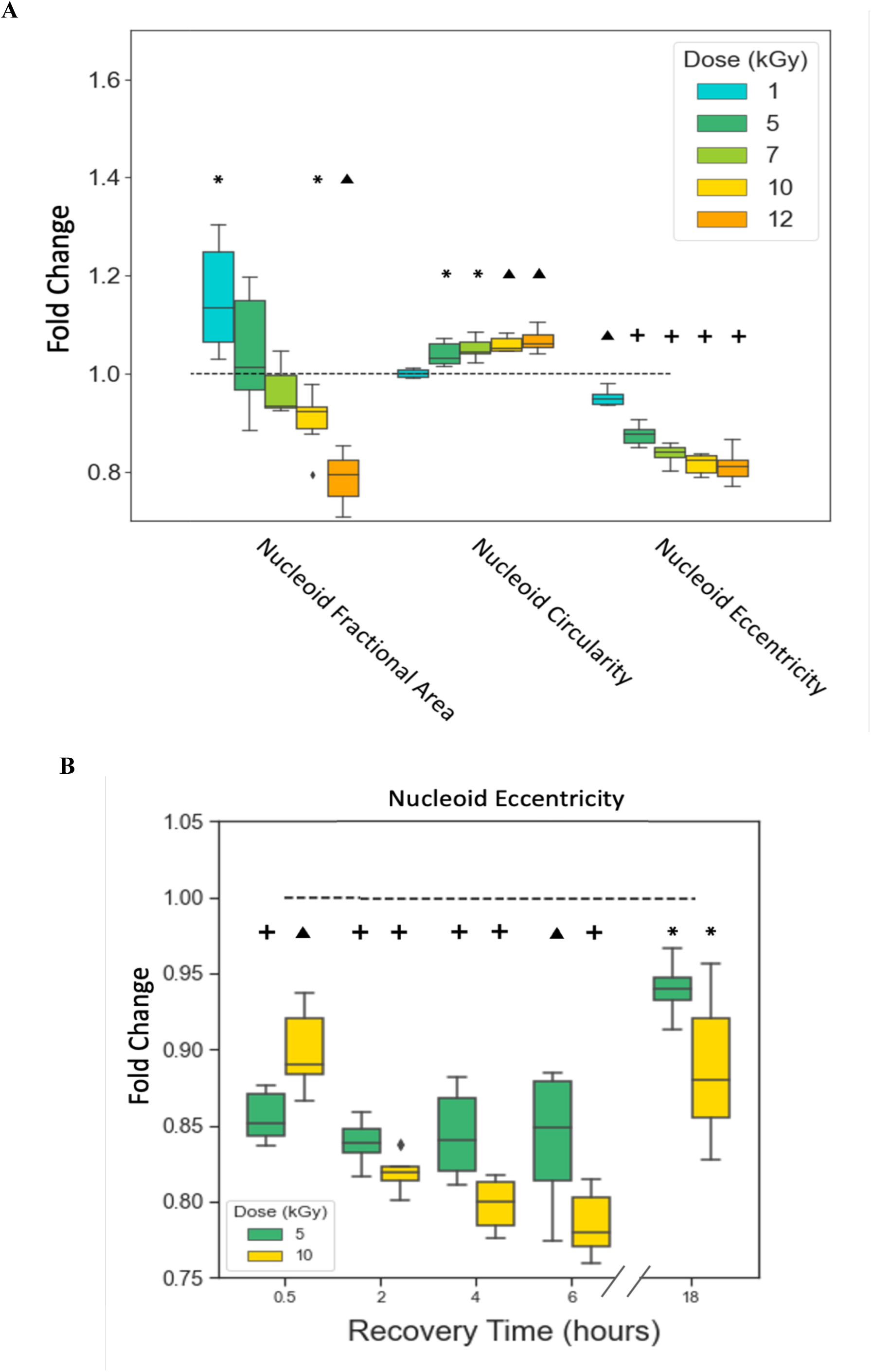
Changes in the geometric characteristics of *D. radiodurans* cells and nucleoids following IR. **(A)** Fold changes of the mean values of geometric properties from the value measured at 0 kGy. Measured images were taken after 2 hours of recovery. Each box-and-whisker plot contains data from 6 biological replicates over 2 experimental days. **(B)** Fold changes of the mean values of eccentricity from the value measured at 0 kGy for the same biological replicate, at the corresponding recovery time point. Each box-and-whisker plot contains data from 6 biological replicates over 2 days of experiments. **(A-B)** P-values are calculated using a student T-test with the null hypothesis that the fold change is equal to 1 (indicated by the black dashed horizontal line). * = p < 0.05, ⏶ = p < 0.001, + = p < 0.0001.

*D. radiodurans* cells exhibited a statistically-significant fold change in both nucleoid fractional area and nucleoid eccentricity at high doses of ionizing radiation. Nucleoid fractional area decreased, nucleoid circularity increased, and nucleoid eccentricity decreased as radiation doses of exposure increased towards 12 kGy (Figure 3A). The decrease in nucleoid fractional area indicates that the nucleoid is becoming smaller with respect to the cell size. The increase in nucleoid circularity indicates that the nucleoid’s perimeter-to-area ratio is decreasing - i.e., it is becoming more condensed. The decrease in eccentricity indicates that the nucleoid is becoming less elongated. Taken altogether, these observations are consistent with the nucleoid becoming more compact.

Next, we measured the dynamics of *D. radiodurans* cell and nucleoid geometries during recovery from ionizing radiation. We monitored unirradiated and irradiated (5 and 10 kGy) *D. radiodurans* populations for 18 hours after exposure to radiation (Figure 3B). Cells were imaged at 0.5, 2, 4, 6, and 18 hours after radiation. The initial four timepoints were chosen to be comparable with the expected doubling time of *D. radiodurans* in our growth conditions (105-130 min) [32, 37]. To measure the continued growth of the cell population, we tracked the post-IR optical density (OD600) of the liquid bacterial cultures over the 18-hour period (Supplementary Figure S1). During this period, we found a steady rate of growth following acute IR exposure at 5 and 10 kGy (Supplementary Figure S1A). This finding suggests that, despite being exposed to significant levels of ionizing radiation stress, *D. radiodurans* cells were able to continue to grow and divide. However, such growth measurements do not reveal the time-course over which cell and nucleoid geometries, having been altered by ionizing radiation, return to the characteristics typifying unirradiated cells.

Next, we explored several important questions; namely, (i) whether all geometric changes observed after irradiation were reversible after a given recovery period, (ii) whether there was a specific time of recovery at which cell and nucleoid changes reverse to pre-radiation geometries, and (iii) whether there was a radiation dose beyond which reversibility in cell and nucleoid changes could not be attained. At each timepoint during recovery, the fold-changes in the geometric properties of each irradiated sample were measured as ratios of mean measured value (for the irradiated sample) to the corresponding mean values of the non-irradiated (control) population of the same biological replicate, at the same time point (Figure 3B, Supplementary Figure S7). The results of significance testing are shown in Supplementary Table S4. We find that recovery after exposure to a 10 kGy IR dose is associated with a greater change in nucleoid eccentricity than is recovery after a lower (5 kGy) IR dose (Figure 3B). After 18 hours of recovery, the fold change in eccentricity for both populations of exposed cells approaches one, indicating no change relative to the non-irradiated cells (characterized by irregular, eccentric shaped nucleoids).

It is worth noting that there are wide distributions of geometric measurements during the 18-hour recovery period (Supplementary Figures S9-13), suggesting heterogeneous sub-populations of cells that may not be properly identified in the bulk data analysis.

### Cluster analysis of single cell morphologies reveals consistent enrichment and depletion of sub-populations with exposure to ionizing radiation

Previous studies have manually identified several classes of nucleoid morphologies in unstressed cells [32]. This led us to computationally identify subpopulations of cells using clustering algorithms. For this analysis, 32 key morphological parameters, encompassing both nuclear and cellular features, were extracted from approximately 470,000 images of individual cells representing all conditions studied. For this analysis, we used a Cell Profiler™ pipeline [38] (Supplementary Table S5). We used a combination of 2D Uniform Manifold Approximation and Projection (UMAP) dimensionality reduction and k-means unsupervised hierarchical clustering to discretize 6 morphological clusters (Supplementary Figure S14) with unique nuclear and cellular morphologies. Importantly, we observed that *D. radiodurans* cells adopt a broad spectrum of morphologies across experimental conditions, with distinct quantifiable spatial and geometric parameters (Figure 4A, Table 1).

**Figure 4.**
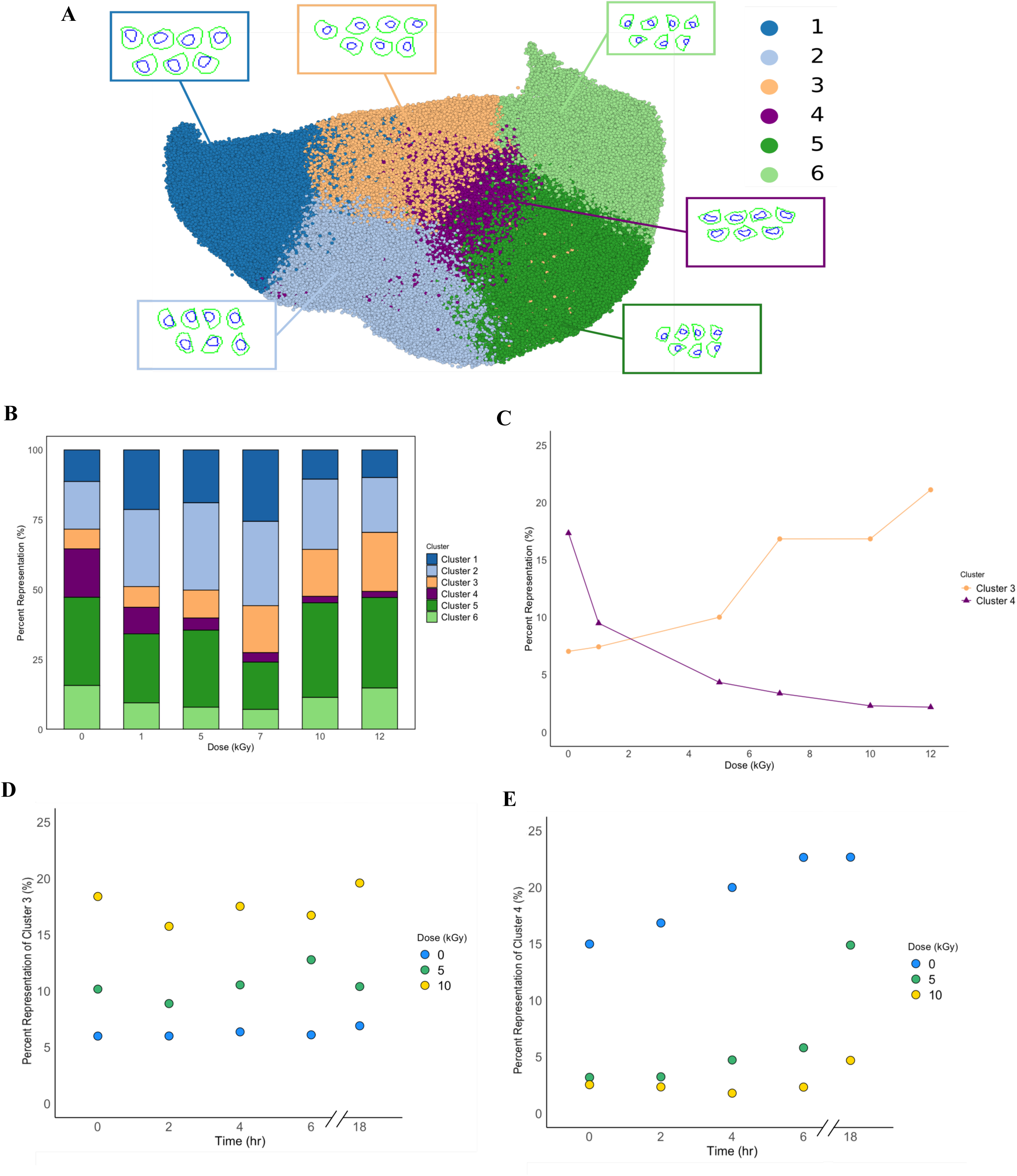
Morphological clustering and analysis of group representation amongst *D. radiodurans* cell populations. **(A)** 2D-Uniform Manifold and Projection (UMAP) construction utilizing the full dataset of collected cell images. Cells are binned into one of six distinct morphological groups. **(B)** Morphological group representation for *D. radiodurans* wild-type strain across acute IR dose treatments. Group representation is displayed as a percentage of the total population of cells contained within each cluster defined in 4A. **(C)** Change in the percent representation of morphological groups 3 and 4 respectively across acute IR dose treatment. **(D)** Change in the percent representation of morphological group 3 over an 18-hour recovery period for 0, 5, and 10 kGy acute IR dose treatments. **(E)** Change in the percent representation of morphological group 4 over an 18-hour recovery period for 0, 5, and 10 kGy acute IR dose treatments. **(C-E)** Lines are drawn as guides to the eye. **(B-E)** Raw percentage values are displayed in k-means enrichment tables (Supplementary Figure S14).

**Table 1.**
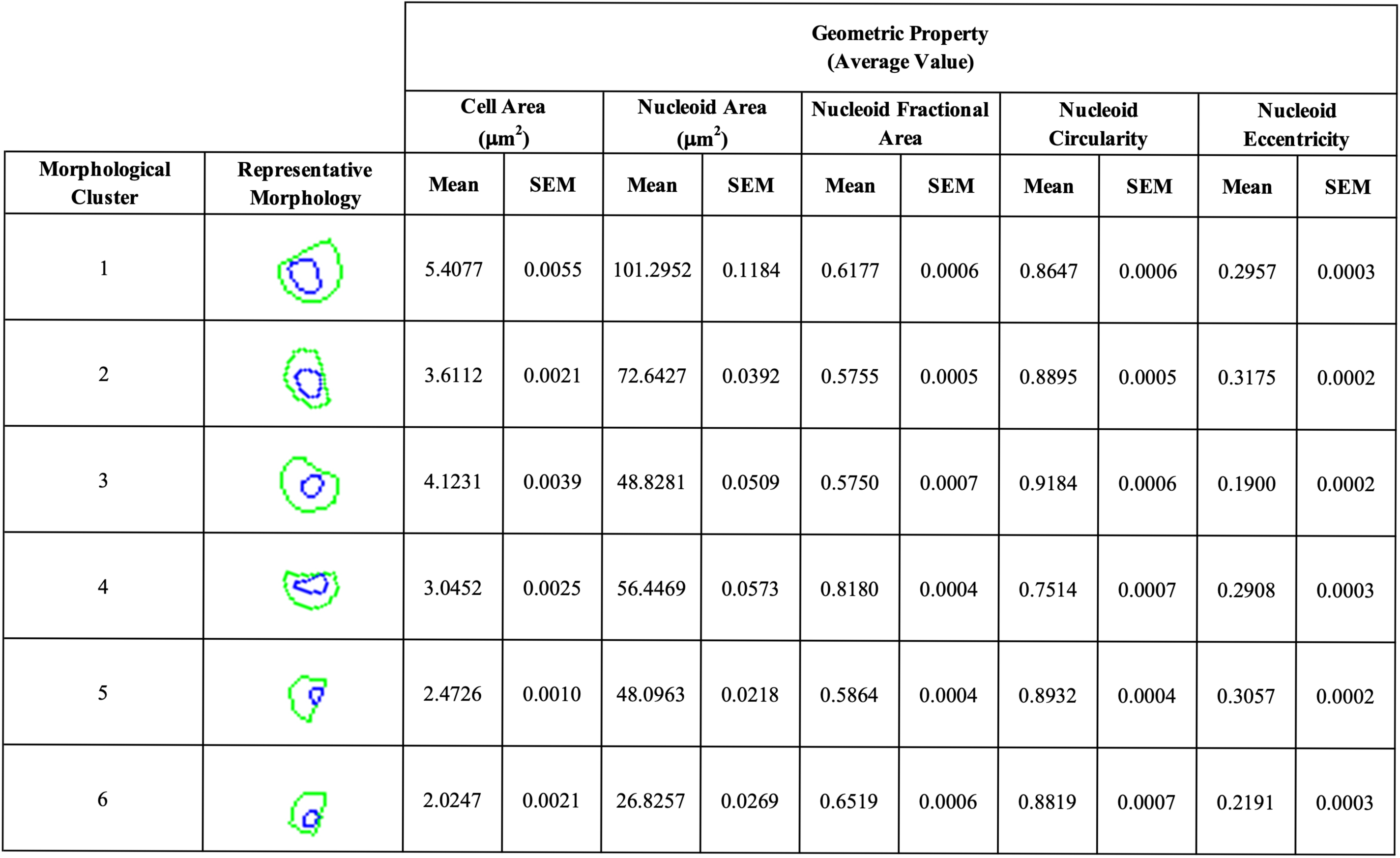
Mean measured geometric properties of morphological groups as defined in Figure 4A. Standard error of the mean (SEM) shown for corresponding geometric property.

| Morphological Cluster | Representative Morphology | Geometric Property<br>(Average Value) |  |  |  |  |  |  |  |  |  |
| --- | --- | --- | --- | --- | --- | --- | --- | --- | --- | --- | --- |
| | | Cell Area<br>($\mu\text{m}^2$) | | Nucleoid Area<br>($\mu\text{m}^2$) | | Nucleoid Fractional Area | | Nucleoid Circularity | | Nucleoid Eccentricity | |
|  |  | Mean | SEM | Mean | SEM | Mean | SEM | Mean | SEM | Mean | SEM |
| 1 |  | 5.4077 | 0.0055 | 101.2952 | 0.1184 | 0.6177 | 0.0006 | 0.8647 | 0.0006 | 0.2957 | 0.0003 |
| 2 |  | 3.6112 | 0.0021 | 72.6427 | 0.0392 | 0.5755 | 0.0005 | 0.8895 | 0.0005 | 0.3175 | 0.0002 |
| 3 |  | 4.1231 | 0.0039 | 48.8281 | 0.0509 | 0.5750 | 0.0007 | 0.9184 | 0.0006 | 0.1900 | 0.0002 |
| 4 |  | 3.0452 | 0.0025 | 56.4469 | 0.0573 | 0.8180 | 0.0004 | 0.7514 | 0.0007 | 0.2908 | 0.0003 |
| 5 |  | 2.4726 | 0.0010 | 48.0963 | 0.0218 | 0.5864 | 0.0004 | 0.8932 | 0.0004 | 0.3057 | 0.0002 |
| 6 |  | 2.0247 | 0.0021 | 26.8257 | 0.0269 | 0.6519 | 0.0006 | 0.8819 | 0.0007 | 0.2191 | 0.0003 |

Clusters 1 and 2 are characterized by enlarged cell areas, enlarged nucleoids, and a large nucleoid fractional area. Cluster 3 has an enlarged cell area similar to that of Clusters 1 and 2 but has a smaller nucleoid area. It is worth noting that Cluster 3 is the most-circular morphological Cluster and has the lowest nucleoid fractional area amongst all Clusters. Cluster 4 has enlarged nucleoids and is the most eccentric geometrically; this morphology has been shown to be associated with active cell division, and therefore perhaps active genome repair [32, 39]. Lastly, Clusters 5 and 6 are characterized by smaller nucleoid areas and nucleoid fractional areas than the other morphological Clusters. Clusters 5 and 6 also have highly circular nucleoids. For non-irradiated *D. radiodurans* cells, Cluster 5 is the most represented morphological Cluster (Figure 4B) while Cluster 3 has the lowest representation.

We suspected that the relative abundance, or enrichment, of these morphological Clusters should be different for cell populations during recovery from exposure to ionizing radiation relative to populations of unirradiated cells. To identify dose-dependent shifts in the enrichment of morphological Clusters, we first analyzed the morphological clustering of cell populations collected in the dose-dependent study of wild-type R1 (Figure 4B). We found that Cluster 3 and Cluster 4 experienced the largest shifts across different doses (Supplementary Figure S14A). During recovery after exposure to a low dose of ionizing radiation (1 kGy), representation of morphological Cluster 4 in wild-type *D. radiodurans* cell populations decreases nearly two-fold. During recovery from irradiation after exposure to higher ionizing radiation doses (10-12 kGy), representation of Cluster 4 decreases nearly eight-fold (Figure 4C). In contrast, representation of morphological Cluster 3 increases over 2-fold during recovery from 10-12 kGy of ionizing radiation (Figure 4C). Representative data for Figure 4C captures recovery after 2 hours. It is worth noting that analysis that upon analysis of the shifts in morphological clusters (Cluster 3 and Cluster 4) across 18 hours of recovery after exposure to ionizing radiation (Supplementary Figure S14B, S15), we observed no further changes in the relative abundance of morphological Cluster 3 (Figure 4D); this was a similar pattern to that observed for the representation of Cluster 4 over time of recovery for the first six hours post irradiation (Figure 4E). However, we observed that the relative abundance of morphological Cluster 4 increased following sham irradiation (0 kGy) in the first 6 hours of recovery, after which Cluster 4 representation seems to reach a steady-state. We also observed an increase in the relative abundance of morphological Cluster 4 from 6 hours post irradiation to 18 hours.

### Screening genomic mutants of D. radiodurans indicate that additional nucleic-acid binding proteins and stress-response proteins affect cell population morphologies during recovery from ionizing radiation

To assess the potential role of different classes of proteins on geometric changes during recovery from ionizing radiation, we used 13 selected genetic knockouts, knockdowns or mutants of genes from 4 distinct classes (Supplementary Table S6). Representatives of the first class, stress-response network genes, included two characterized regulatory stress response proteins (pprA, IrrE/pprI) and one characterized stress response regulatory small RNA (Dsr2/PprS) [26]. Representatives of the second class, nucleic-acid binding proteins, included four proteins that (in other organisms) associate with small RNAs under stress (KphA, KphB, PNPase, and Rsr). The third class is composed of proteins characterized in *D. radiodurans* that have homology to known *E. coli* nucleoid-associated proteins (NAPs). These proteins include two *E.coli* NAPs (Dps, RecA) and a deletion of the 5’ regulatory untranslated region encoding for GyrA (-gyrA, another predicted NAP involved in DNA repair) (Villa et al 2017). The fourth and final class is represented by known and established *D. radiodurans* NAPs (Lrp, DR_1116 and DRA_0141) [30]. These 13 genomic knockouts in four categorical classes were selected from a pre-existing library or generated for this publication (Supplementary Table S6-S7).

We first characterized geometric properties of the protein mutant strains, in the absence of radiation, and compared these properties to those of the wild-type R1 strain. Measured changes in cell area, nucleoid area, nucleoid fractional area, nucleoid circularity, and nucleoid eccentricity are shown in Supplementary Figure S16. Mutant strains had average cell areas very similar to the wild-type, with no differences greater than 20% (although some were statistically significant). In contrast, the nucleoid area and the nucleoid fractional area were at least ∼ 20% greater in at least one of the known *D. radiodurans* NAP (DRA_0141) and another hypothesized *D. radiodurans* NAP (Dps). This finding validates our ability to capture the basal role of these proteins in maintaining nucleoid size, even in the absence of any radiation stress. We also found a potential, not yet reported, role of KhpA and KhpB in maintenance of nucleoid size under unstressed conditions, in the form of a ∼20% difference in the nucleoid area of the genomic knockout strains for these proteins. We found only minor (although statistically significant) differences of less than 10% in nucleoid eccentricity and circularity in all the mutant strains relative to the wild-type strain under these unstressed conditions.

To identify associations between specific proteins and response to radiation, we exposed each mutant strain to sham (no radiation) and to 10 kGy of IR. For these experiments, the wild-type strain was also simultaneously irradiated and re-characterized as an internal experimental control. We measure the relative abundance of the six morphological Clusters defined above (Figure 5A) after two hours of recovery from IR or sham (Figure 5, Supplementary Figure S17). We find eight strains, *ΔDRA0141*, *ΔGyrA-UTR, ΔLrp, ΔDR1116, ΔpprI, ΔKhpA, ΔKhpB,* and *ΔRsr,* which have at least one morphological Cluster with a percent representation that is different from that of the wild-type by at least a factor of two at the same dose of ionizing radiation (Figure 5A-D, Supplementary Table S8). We quantify this for each morphological Cluster using a logarithmic scale, defined as: *log2FoldChange = log2(percent representation in mutant/ percent representation in wild-type).* A factor of two differences in representation corresponds to log2FoldChange (log2FC) ≥ 1 or ≤ -1. Among these eight strains, the most consistent changes, defined as a p-adjusted (p-adj) value less than 0.05, were found in six strains: *ΔDRA0141*, *ΔGyrA-UTR, ΔLrp, ΔDR1116, ΔpprI,* and *ΔRsr* (Figure 5A-D, Supplementary Table S8).

**Figure 5.**
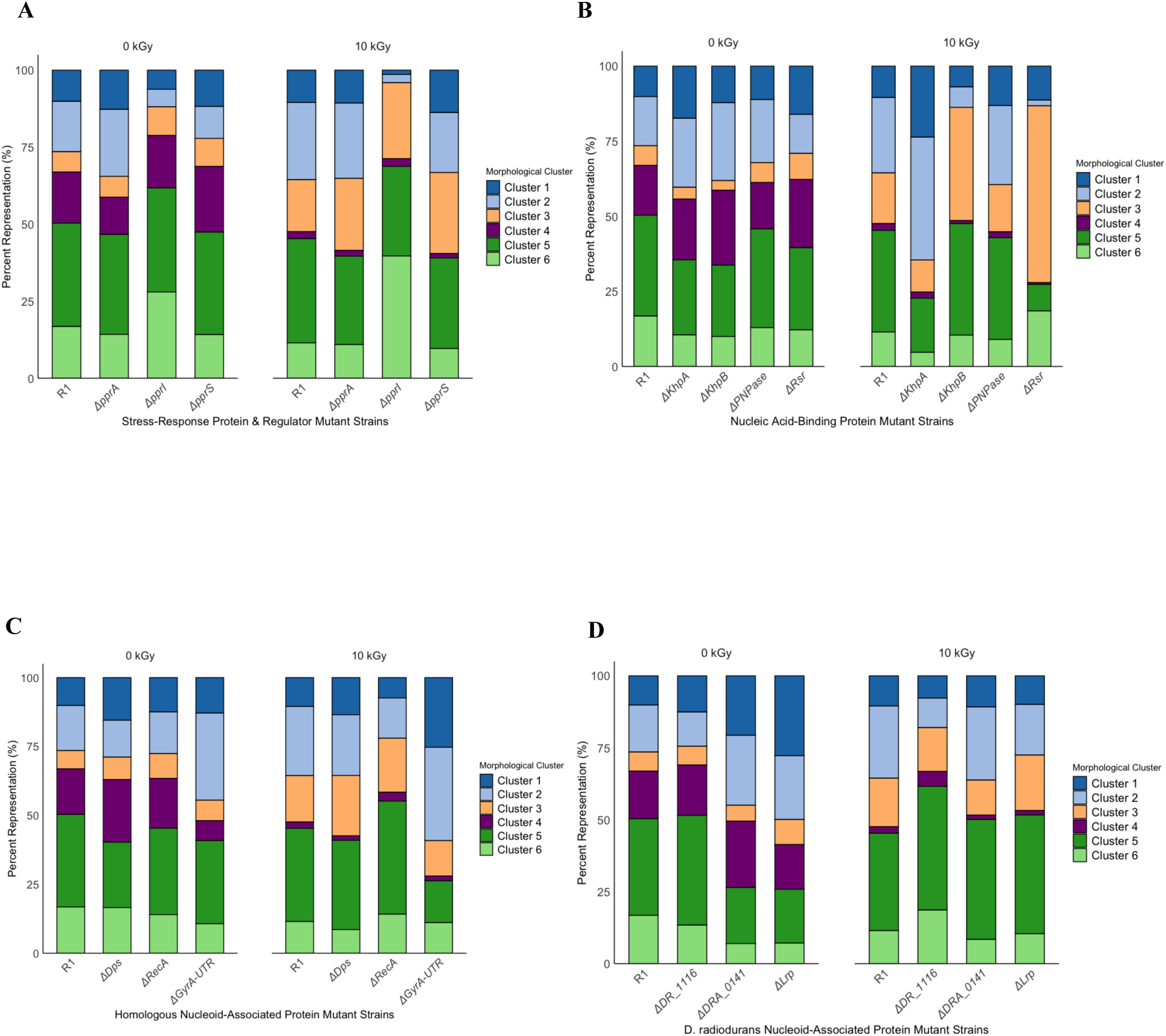
Shifts in morphological group representation with acute IR dose treatments for *D. radiodurans* genomic mutant strain classes. Group representation is displayed as a percentage of the total population of cells contained within each cluster defined in Figure 4A. **(A)** Stress-response proteins & regulator. **(B)** Nucleic acid-binding proteins. **(C)** *D. radiodurans* proteins with homology to *E. coli* nucleoid-associated proteins. **(D)** Known *D. radiodurans* nucleoid-associated proteins.

## Discussion

There are growing databases of newly discovered organisms and bacterial communities that include data sets of genomic, proteomic, and transcriptomic information [40–43]. Efforts have been made to predict bacterial phenotypes from this information [44]. However, dynamic phenotypic changes in nucleoid organization have not been well characterized in extremophilic organisms undergoing stress. The connection between oxidative stressors (i.e. ionizing radiation) and mechanisms of DNA protection and repair continues to be of interest, particularly in light of genome maintenance by extremophilic organisms. Unlike proteobacteria such as *E. coli*, *D. radiodurans*, a model extremophile known for its unique stress tolerance and genome maintenance mechanisms [45–48], constitutively maintains a highly-condensed nucleoid. Thus far, quantitative characterization of nucleoid geometric features and their dynamics under stress have not been reported in *D. radiodurans*.

To address this gap, we have developed a high-throughput pipeline for image analysis and used it to characterize geometric properties of nucleoids of the extremophile bacterium *D. radiodurans*. The user-defined geometric properties that we measure are the cross-sectional area of cells and nucleoids, the ratio of nucleoid cross-sectional area to cell cross-sectional area (which we term “nucleoid fractional area”), and the circularity and eccentricity of nucleoids. We find that IR results in significant changes in the geometric properties of wild-type R1 cells. Specifically, the average cell area is greater following IR, and the average nucleoid eccentricity is lower, for all radiation doses tested (Figure 3A, Supplementary Figure S7, Table 1). The increase in cell area is consistent with the idea that bacterial cells under stress divide less frequently and therefore may grow to larger sizes before division [34, 49]. The changes in nucleoid eccentricity and circularity both point to the nucleoid becoming a more-compact and more-spherical shape after radiative stress than before. Moreover, these changes in nucleoid geometry both show monotonic trends with radiation dose, which may suggest a dose-dependent effect of IR on increasing compaction and sphericity in the *D. radiodurans* nucleoid (Figure 3A, Supplementary Figure S7). Therefore, we conclude from our work that changes in geometric measurements of morphology during an extended time after acute radiation exposures might reflect recovery from IR stress. Recovery from IR stress is of particular interest in the case of *D. radiodurans* because it has the potential to yield insight into this extremophile’s unusually-high tolerance of IR.

While bulk-data analysis of mean properties of the whole, consolidated population yields general insight into the shifts in geometric properties in *D. radiodurans*, the probability-density distributions of these measurements are broad (Supplementary Figure S2-S6). This represents high variance in the data and suggests that the geometric properties found in these populations may be heterogeneous. To expand the insight yielded by image analysis beyond that achieved by user-defined characteristics, we used a computational modeling algorithm, on a parameter space defined by 32 different morphological characteristics. This algorithm clustered the 400,000+ imaged cells into six different morphological Clusters within the 32-dimensional parameter space. One of these Clusters, Cluster 4, is characterized by elongated nucleoid shapes (Figure 4A, Table 1). Previous work has shown that elongated nucleoids are associated with cell division [32, 39] and are notably less present in bacterial populations following acute radiation [29]. Thus, the depletion of morphological Cluster 4 following IR stress agrees with what is known about bacterial biology in general and also yields new insight into the response of *D. radiodurans* specifically to IR stress. In contrast, Cluster 3 representation is enriched by IR in a dose-dependent manner. Morphologically, Cluster 3 has the most circular nucleoid and the smallest nucleoid fractional area amongst the defined Clusters (Table 1). Thus, Cluster 3 has the most compact nucleoid shape. The dose-dependence of Cluster 3 enrichment suggests that these geometric properties may indicate that DNA repair is aided by coalesced genomic material. These clear trends, with increasing IR dose, of increasing enrichment of Cluster 3 and increasing depletion of Cluster 4, appear to be signatures of recovery following radiation exposure (Figure 4B).

Following survival comes recovery, which is a time-dependent process. *D. radiodurans’* recovery from IR-caused stress is a 3-step process of DNA degradation, DNA recombination, and a resumption of DNA synthesis [45, 50]. DNA degradation and delay in DNA synthesis have been shown to be dependent on radiation dose [20]. Indeed, our morphological analysis shows that the timescale for recovery appears dependent on the radiation dose (Supplementary Figure S1). We have quantified the geometric properties of *D. radiodurans* populations over the 18 hours following IR at an intermediate dose (5 kGy) and a high near-lethal dose of 10 kGy (which is approaching the D10 value of 12 kGy).

Our morphological analysis identified sub-population behavior not only in a dose-dependent manner, but along the *D. radiodurans* recovery timescale. Following IR dosing at 5 or 10 kGy, Cluster 3 representation is roughly constant over the next 18 hours of recovery (Figure 4C). Thus, even hours after radiation exposure *D. radiodurans* cells have compacted nucleoids, as indicated by the low nucleoid fractional area and highly circular nucleoid characterizing morphological Cluster 3. We speculate that this morphological Cluster could act as an identifiable long-term marker of DNA damage induced by IR.

If the geometric changes observed following radiation are linked to the activity of the molecular machinery for DNA repair and recovery, it is plausible that changes in this machinery may be reflected in *D. radiodurans*’ structural characteristics even in the absence of radiation stress. Prior studies have identified several gene products that may impact nucleoid organization and structure [30, 51]. We evaluated 13 selective genomic mutant strains with deletions of or mutations to genes that encode for proteins involved in stress response, binding to DNA or RNA, and nucleoid association (Supplementary Table S6). For every mutant strain tested, at least one user-defined geometric property was significantly different from the corresponding property in the wild-type (Supplementary Figure S16). These data likely suggest that the morphological clusters will differ in the absence of these gene products when compared to wild-type.

Computational morphological clustering has the potential to yield insight into the roles of specific gene products both in *D. radiodurans’* resilience to radiative stress and in regard to this organism’s constitutively-condensed nucleoid. Knocking out such gene products results in several cases where the population of these morphologically-extreme Clusters is shifted from that of the wild-type R1. These shifts can be seen in the changes in representation by morphologically-extreme Clusters upon gene deletion at sham conditions (Table 1). These morphologically-extreme Clusters are Clusters 1, 2, and 6. Clusters 1 and 2 are characterized by a large nucleoid and large nucleoid fractional area; Cluster 6 has a small nucleoid and small nucleoid fractional area. Notably, these shifts were often in accord across Clusters - in such cases, Clusters 1 and 2 increased or decreased together, and Cluster 6 decreased or increased inversely (Figure 6). However, in other cases Cluster 2 increased while Cluster 1 was largely unchanged. The contrast between correlated and anti-correlated shifts for Clusters 1 and 2 show the potential of this approach to yield insight into different molecular mechanisms underlying the maintenance of *D. radiodurans’* constitutively-condensed nucleoid.

If genomic mutants respond to IR similarly to wild type *D. radiodurans*, we expect to see enrichment of morphological Clusters 3 and depletion of morphological Cluster 4 as characteristic of the recovery period after exposure to IR (Figure 4B). Indeed, all mutant strains tested experience a depletion in morphological Cluster 4 after IR exposure (Figure 5A-D, Supplementary Table S8). However, for two strains (KhpB and Rsr), we find that morphological Cluster 4 is even more enriched (log2FoldChange > +/- 1) after IR than it is for the wild-type; both KhpB and Rsr are classed as nucleic-acid binding proteins. Additionally, we find that the mutant lacking the stress response protein PprI has a statistically-significant, log2FoldChange > 1, greater representation than the wild-type in the condensed morphological extreme, Cluster 6, after radiation. In all three cases (mutants lacking KhpB, Rsr, or PprI), we see lower representation (inverse log2Fold) of of the decondensed extreme, Cluster 2 after the radiation than for the wild-type after radiation (Rsr and PprI are statistically significant - p.adj < 0.05). This suggests that removal of any of KhpB, Rsr, or PprI from the *D. radiodurans* genome results in a greater nucleoid condensation upon exposure to IR stress.

In contrast with the three single-protein mutations that result in greater condensation following IR, another member of the nucleic acid-binding class (KhpA) has an opposing effect. The KhpA mutant has greater representation, log2FoldChange > 1, of Cluster 1 following IR than does the wild-type; Cluster 1 is a decondensed-nucleoid morphological extreme. Following IR, the KhpA mutant less representation, log2FoldChange < -1, of Cluster 6 than does the wild-type; Cluster 6 is a condensed-nucleoid morphological extreme, Cluster 6.

Surprisingly, deletions in proteins previously described as nucleoid-associated in *E. coli* or *D. radiodurans* were much less impactful on morphology than were deletions in stress-response and nucleic acid binding proteins. Of the nucleoid-associated proteins studied, only one, DR_1116, differed from the wild-type by log2Fold in morphological representation following IR. Rather, the nucleic acid-binding and stress response class of proteins observed in this study, specifically KhpA, KhpB, Rsr and PprI, appear to have the most notable effects on *D. radiodurans* nucleoid morphology following IR. Importantly, these proteins were not previously known to play a role in controlling nucleoid dynamics and condensation in bacterial populations. This suggests that proteins from these two known classes in *D. radiodurans* may play critical, undescribed roles in dynamic nucleoid regulation, meriting further study.

Interestingly, *ΔGyrA-UTR* exhibited a phenotype under IR stress of various cell membrane protrusions (Supplementary Figure S18). This phenotype appears similar to blebbing behavior, typically seen in mammalian cells under apoptosis, stem-cell differentiation, and other “cell-fate” determination states [52]. While quantitative determination of blebbing for the purpose of tracking signaling pathway activation has occurred in mammalian cells [53–55], this method of spatiotemporal organization analysis has not yet been utilized in bacteria. These results warrant further exploration as they may provide insight into potential signaling pathways in *Deinococcus* under IR stress.

We expect that this method we present here should be able to be used in the analysis of other prokaryotes to measure the geometric properties of cells and nucleoids. For the nucleoid in particular, geometric parameters such as compactness and size impact biological processes essential to genome repair through mechanisms such as restricted diffusion and crowding [56, 57]. The approach we present here can be used to measure changes in nucleoid compaction, as well as other morphological changes, across different organisms to provide new insight into their stress response mechanisms and their survival. It also has the potential to be combined with more-standard genetics techniques to yield insight into the regulation of subcellular compartmentalization in a wide range of bacteria.

Furthermore, the analysis techniques used here are not specific to bacteria; this approach should be directly extensible to a wide range of organism types and to types of subcellular structures that are not nucleoids or DNA-containing. Given differences across organisms in chromosomal length, copy numbers, and survivability across different types of environmental stressors, the connection between the dynamics of nucleoid dynamics organization and the presence of extreme phenotypic traits in bacteria remains intriguing. Geometric parameters such as compactness and size impact biological processes essential to genome repair such as restricted diffusion and crowding (Minsky 2003, Esadze and Stivers 2018), thus quantifying nucleoid compaction activity across different organisms is a key aspect of understanding their stress response mechanisms and their survival. Therefore, the work presented here can be used to probe morphological features at the subcellular level and expose biophysical phenomena not readily recognizable otherwise.

The utility of this method is likely to be increased by the use of high-resolution and super-resolution imaging techniques, which will make finer gradations in geometry distinguishable and may allow the possibility of examining an even higher-dimensional morphological space.

## Conclusion

We created a robust image analysis methodology to elucidate quantitative features of bacterial cell and nucleoid morphology. These methods allowed quantitative analysis of geometric changes in the cell envelope structure and the nucleoid structure of the extremophile bacteria *D. radiodurans* following ionizing radiation and following genomic deletion of key candidate proteins. We find dose- and time-dependent changes in the cell and nucleoid morphologies. The methods create a useful screening tool to probe subcellular organization and its regulation, response to stress, and possible mechanistic roles of genome-associated proteins in nucleoid condensation.

## Methods

### Bacterial cultures and growth conditions

*Escherichia coli* strain DH10b and its derivatives were grown at 37°C in Luria-Bertani (LB) media broth (10 g/L tryptone, 10 g/L NaCl, and 5 g/L yeast extract) or LB solid (1.5% agar) medium. When necessary, antibiotics were used at a concentration of 34 µg/ml (Chloramphenicol) for *E. coli* strains. *Deinococcus radiodurans* strain R1 (ATCC 13939) and its derivatives were grown at 32°C in TGY broth (1% tryptone, 0.1% glucose, 0.5% yeast extract) or TGY solid (1.5% agar) medium. When necessary, antibiotics were used at a concentration of 16 µg/ml (Kanamycin, Tetracycline), 3.4 µg/ml (Chloramphenicol) for *D. radiodurans* strains. All strains are listed in Supplementary Table S6.

### Transformation of recombinant plasmids in *D. radiodurans*

Recombinant plasmids were transformed into *D. radiodurans* as previously described [51]. *D. radiodurans* cells were grown to late log phase (OD600 = 1.0) and mixed with 30 mM CaCl_2_ and 10% glycerol. For transformation into the competent cell stock, 1.5µg Plasmid DNA was added, followed by an incubation of 1 hour on ice, and a subsequent incubation at 32°C for 1 hour. Cells were then incubated at 32°C shaking overnight in 800µl of fresh TGY media in test tubes. Post-incubation, cells were plated onto TGY plates with the appropriate antibiotic concentration (above), and incubated for 3 days at 32°C. Colonies were verified via polymerase chain-reaction (PCR) and sequencing (SNPsaurus).

### Construction of genomic knockout strains in *D. radiodurans*

The knockout strains of *D. radiodurans* were constructed using a homologous recombination method reported previously [27]. In short, 1kb regions upstream and downstream from the gene selected for deletion were amplified via PCR from the R1 genome, and cloned into the pUC19mPhes plasmid alongside a fragment containing a kanamycin cassette, as well as lox66 and lox71 sequences. All fragments were then gel-purified using QIAquick Gel Extraction Kit (Qiagen) and assembled with HindIII-digested pUC19mPheS plasmid using NEBuilder® HiFi DNA Assembly Master Mix (New England Biolabs Inc.). The resulting plasmid was then transformed into *E.*c*oli* DH10b strain for isolation via QIAprep Plasmid Mini-Prep Kit (Qiagen). The isolated plasmid was subsequently transformed into *D.radiodurans* R1 chemically competent cells (above), and underwent double-crossover homologous recombination. Mutant strains were selected on TGY plates with the appropriate kanamycin concentration and 4-chlorophenylalanine (5 mM) for 3-5 rounds of selection. The total process selected for cells that underwent homologous recombination to replace the target region with the kanamycin cassette, and no longer expressed the pUC19mPheS recombinant plasmid. The kanamycin resistance marker was removed using Cre/Lox recombination, after transformation of the pDeinoCre plasmid into competent cells of the deletion strain (above). The primers used for these processes are included in Supplementary Table S7. All plasmids are listed in Supplementary Table S6.

### Exposure of bacteria to ionizing radiation (IR)

For acute IR exposures, exponential phase cells (OD600 = 0.8) were irradiated with a 10-MeV, 18-kW linear accelerator (LINAC) β-ray source at dosages ranging from 0 to 12 kGy (250 Gy/s) at the National Center for Electron Beam Research, Texas A&M University, as reported previously [26, 51, 58]. Samples were kept in stasis on dry ice during transport to and from the irradiation facility (∼2 h each way). Irradiated samples were then incubated in TGY broth for no additional recovery time (for survival assays and growth assays) or 2 h (for staining and imaging) at 32 °C immediately following irradiation and transport. Exact acute IR dose exposures and dose ranges for each day of experimentation are located in Supplementary Table S10.

### Survival assays during recovery post-IR

To measure survival following acute IR exposures, biological replicas in triplicate of exponential phase cells (OD600 = 0.8) were irradiated with a 10-MeV, 18-kW linear accelerator (LINAC) β-ray source at doses 0, 1, 5, 7, 10, 12 kGy (250 Gy/s) at the National Center for Electron Beam Research, Texas A&M University as reported previously [26, 51, 58]. Following irradiation at room-temperature and transport, samples were immediately serially diluted (10 ^-^0 to 10 ^-^3) and plated onto TGY agar plates. Plates were incubated at 32 C° for 2 days, and colonies were counted. Relative survival rates were defined as the percentage of colony forming units observed under each IR dose condition compared to samples that received no IR.

### Growth assays during recovery post-IR

Growth curves of R1 and RecA KD at variable kGy dosages were evaluated using a Plate Reader (BioTek) and OD_600_ was calculated using a bacterial calibration curve for *D.radiodurans* strains. Following irradiation at room-temperature and transport, samples were immediately incubated in culture tubes at 32 C°. Biological replicates of each strain (25 µl), at each dosage interval, in technical triplicate, were distributed into 96-well plates (Greiner Bio-One µClear™ Bottom 96-well Polystyrene Microplates) with 175 µl TGY media. OD_600_ of each sample was measured every 2 h for 18 h as the cultures grew with shaking at 32°C, and the average of all replicas per strain, per dose condition, was calculated.

### Staining Protocol

Cells were stained using 3 mM stock solution of Nile Red (Thermo Fisher, Invitrogen) (used for staining membranes) and 15uM stock of Syto 9 green nucleic acid stain (Thermo Fisher, Invitrogen) (used for staining nucleoids). Stock solutions of Nile Red were made by dissolving solid Nile Red stain into DMSO at a concentration of 3mM. Stock solutions of Syto 9 were diluted to 15uM from factory stock into DMSO. In 1 mL of cell culture, final concentrations of Nile Red and Syto 9 were 30 uM and 150 nM respectively. Cells were then placed in a rotary shaker for 30 minutes at 32°C. Cells were then spun down and stain solution was removed. The cell pellet was then resuspended in 200 uL of Dulbecco’s Phosphate Buffer Solution (DPBS). Cell suspensions were then prepared for imaging by putting 15 uL of cell solution onto a microscope slide with an image spacer (Grace Bio-Labs) and a #1 size coverslip (Fisher Scientific).

### Imaging

Microscopy is done via confocal microscopy using a Fluoview 1000 confocal microscope on an Olympus IX81 microscope base. The 488 nm and 543 nm laser lines were used. The Fluoview settings are described in Table 2.

**Table 2.** Confocal Microscope software settings. PMT voltage is provided as a range due to voltage variation required for a low saturation of pixels. Voltage selection is performed via adjustment of the Hi-Lo LUT edit setting and changing the voltage to provide a minimal number of red pixels while maximizing grayscale pixels

| Laser (nm) | Power (%) | PMT Voltage (V) | Gain | Offset (%) |
| --- | --- | --- | --- | --- |
| 488 | 5 | 350-450 | 1x | 20 |
| 543 | 15 | 600-700 | 1.5x | 20 |

### Image analysis protocol

Z-stack images are opened in FIJI, which is an open-source, free image analysis software [59]. The z-slice that is at the midpoint of the cells are found by iterating through the z stack and picking the midpoint by eye. This slice is then saved into 2 parts, a membrane image and a nucleoid image. These images are saved with the same name as the z stack with the suffix “_mem.tif” and “_nuc.tif” respectively. The rest of the analysis is done in a python (https://ipython.org) jupyter notebook [60, 61]. The notebook opens the images and creates a composite image with one channel being the membrane image and the other being the nucleoid image. These composite images are then segmented using the cellpose application programming interface (API) [33]. The nucleoids are segmented using the ‘nuclei’ model in cellpose with the initial diameter set to 9 pixels and flow_threshold equal to 0.4. The cells are segmented using the ‘cyto’ model using both the membrane channel and the nucleoid channel, with initial diameter set to 13 pixels and flow_threshold equal to 0.4. The masks from each model evaluation are saved with the suffix ‘_nuclei_mask.tif’ and ‘_cell_mask.tif’ respectively. These masks then go through a process of connecting the nuclei and cell masks. The connecting process does the following: ensure each nucleoid and cell are matched 1:1 with the same pixel value, ensure no 2 nuclei are within a cell, and ensure nuclei are not bigger than the cell. These masks are then used to get the region properties of each cell/nucleoid pair. The physical quantities measured are nucleoid circularity, nucleoid eccentricity, nucleoid area, cell area, and nucleoid fractional area. Total cells imaged in these analyses for all experiments are located in Supplementary Table S11. Code for the analyses have been made available in a GitHub repository (github.com/bniese/drad-image-analysis).

### Calculation of Geometric Properties

The area and perimeter of each nucleoid mask can be used to find its circularity, as follows: 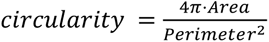. Circularity measures the degree to which the geometry of the nucleoid mask approaches a mathematically perfect circle, for ; *Area* = *πr*^2^ and *Perimeter* = *πr*^2^, where *r* is the radius of the circle. Circularity has a value of one for a perfect circle, which is the shape that maximizes the area-to-perimeter ratio. Thus, as the nucleoid becomes more compact, the circularity will increase. Artifacts in our analyses of digital images of nucleoids can give circularity values greater than one, as follows: The digital nucleoid mask is made of square pixels. The perimeters of the nucleoid masks are estimated using a built-in function in the scikit image library in Python. Parts of the mask pixels can lie outside the bounds of the estimated perimeter. If enough mask area is located outside the estimated parameter, a circularity value greater than one will be calculated. This artifact will consistently act to shift calculated circularity values higher than the true value. However, the consistent direction of this shift means that circularity as used here is still a valid measure of the relative compactness of nucleoids.

Eccentricity is measured by fitting an ellipse to the outside edge of the nucleoid mask. A non-circular ellipse has two perpendicular axes with different lengths, *a* > *b*. Eccentricity < arises from the difference in these two lengths, thus: 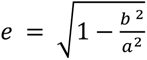. For a circle, *a* = *b*. As the nucleoid becomes less elongated, eccentricity will decrease toward zero (Figure 2B, right side).

### Statistics

Statistical analysis was done in python using the scipy stats library. Kolmogorov-Smirnov test was calculated using the kstest function. Student t-test was calculated with the ttest_1samp function. ANOVA test was calculated with f_oneway function. Number of cells per biological replicate can be found in Supplementary Table S11. For each of the morphological Clusters, pair-wise comparisons of relative abundance, using a t-test, were made between the R1 data and the mutant strains. For example, the relative abundance of Cluster 1 across the R1 samples was compared against the relative abundance of Cluster 1 across the *ΔRecA* samples. A Bonferroni correction was applied to the resulting p-values to correct for the large number of comparisons. Clusters from mutant strains with a change in a relative abundance +/- Log2 of 1 and with an adjusted p-value below 0.05 are reported as significantly different than R1 (Supplementary Table S9).

### Morphological Analysis

A total of 32 key morphological parameters encompassing both nuclear and cellular features were extracted from individual cells using a Cell Profiler^TM^ pipeline [38] (Supplementary Table S5). The optimal number of clusters were determined using the ‘elbow method’, or the beginning of the elbow of the inertia/silhouette graph shown in Supplementary Figure S19. Cluster partitioning was determined by the k-means algorithm, an unsupervised approach that only requires the number of clusters to be input by the user [62, 63]. Approximately 470,000 cells encompassing all experimental conditions were used to capture the entirety of possible morphological phenotypes. To standardize the dataset, morphological parameters were log-normalized and standard scaled (to account for parameters of different orders of magnitude). The dataset was then used to construct a 2D-Uniform Manifold and Projection (UMAP) space [64]. UMAP is a nonlinear dimensionality reduction algorithm that seeks to capture the structure of high dimensional data in a lower-dimensionality space (for this work, the 32-vector space was simplified to two). Each point in the space represents an individual cell whose morphological properties have been projected. As a synergistic approach to UMAP, k-means clustering, an unsupervised hierarchical clustering algorithm, was performed on the normalized dataset to discretize unique morphological Clusters.

The algorithm converged on six optimal clusters based on a plateau of inertia and silhouette values (Supplementary Figure S19), and these clusters were projected onto the UMAP space for visualization. Together, each individual cell expressed a unique position on the UMAP space and belonged to a k-means morphological Cluster. Condition-specific morphological quantification was determined by the ensemble effects of individual cells within that condition. Given the quantity of cells for this analysis, kernel density estimation with gaussian priors were used to construct contour maps to identify “hot zones” in the UMAP for certain conditions.

## Supporting information

Supplemental Data Sheet 1

Supplemental Data Sheet 2

## Data Availability Statement

The authors confirm that the data supporting the findings of this study are available within the article and its supplementary materials. The code used to generate data sets from microscopy images can be found at github.com/bniese/drad-image-analysis.

## Acknowledgements

We thank the Contreras Group for assistance with overnight *D. radiodurans* growth curves and radiation experiments, especially Trevor Simmons. We would like to thank Gina Partipilo for assistance in developing standardized curves for our plate reader assays. We also thank Bryan Dinh for fluorescence microscopy assistance. We additionally thank Jazmine Johnson and Jennah Johnson for their assistance in culturing *Deinococcus radiodurans* strains for this work. We also thank Dr. Pascale Servant (Université Paris-Saclay), Dr. Michael Daly (Uniformed Services University), and Dr. Suzanne Sommer (Institut de Génétique et Microbiologie - Université Paris-Sud) for their generosity in previously providing us with the following *Deinococcus* mutants used in this study – *ΔDps* (GY13314), *ΔpprI* (GY14127), and *ΔRecA* (recA30 :: kan). Figures 1A and 2B were created with BioRender.com.

