## Supplemental Data Sheet 2 for "Geometric changes in the nucleoids of Deinococcus radiodurans reveal involvement of new proteins in recovery from ionizing radiation"

**A.**

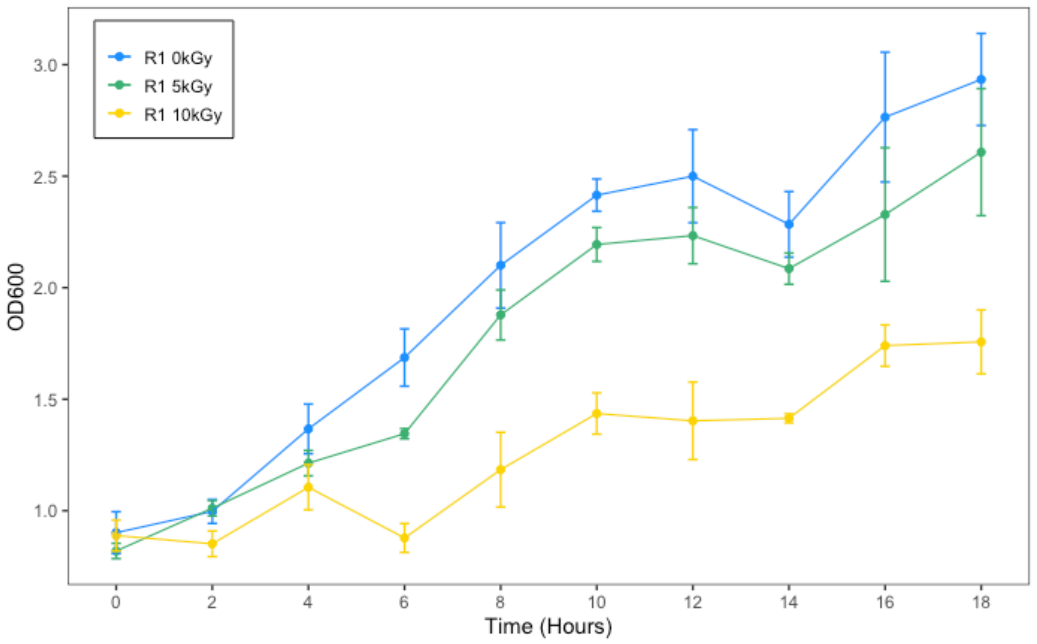


**B.**

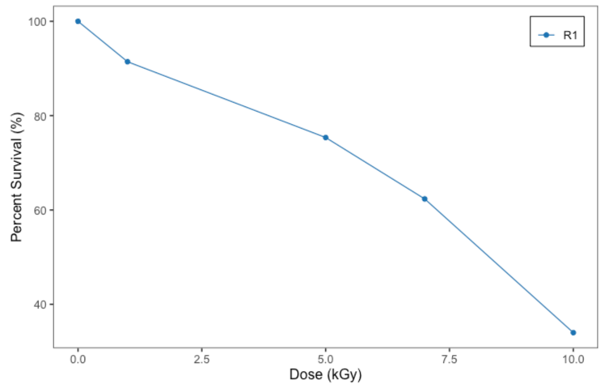


**C.**

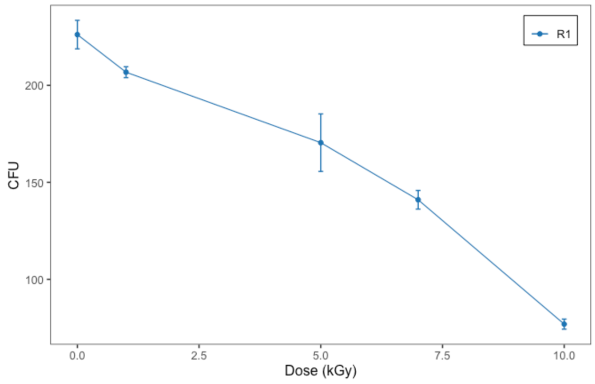


**Supplementary Figure S1.** Growth and survival data for *D. radiodurans* wild-type cells under IR stress. **(A)** Growth curves of the wild-type strain of *D. radiodurans* after exposure to 0, 5, or 10 kGy acute IR. Radiation dose is indicated by the color of the line, as indicated in the legend. Lines represent the average absorbance at $OD_{600}$, and error bars represent the standard error of the mean of technical triplicate of three biological samples (*N* = 9). **(B)** Survival of the wild-type strain of *D. radiodurans* relative to the survival of the bacteria's exposure to 0 kGy of radiation. **(C)** Mean survival of the wild-type strain of D. radiodurans (Plotted as Colony Forming Units [CFU]) as a function of dose (0, 1, 5, 7, 10 kGy). Error bars represent standard deviation of technical triplicate of triplicate biological samples (*N* = 9).

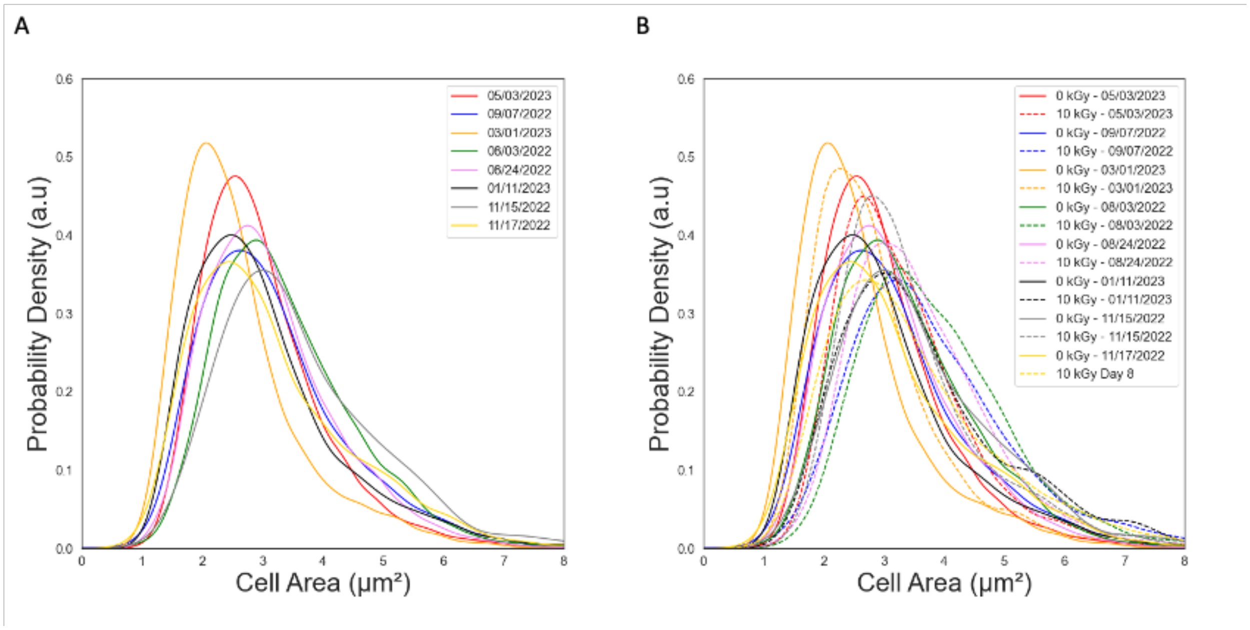


**Supplementary Figure S2.** Probability distributions of cell areas for eight experiments each composed of 3 biological replicates, measured on the dates indicated in the legend. (A,B) For each biological replicate, the probability distribution of cell areas for the non-irradiated (control) sample is shown with a solid line, and (B) the probability distribution of cell areas for the sample irradiated at 10 kGy is shown with a dashed line of the same color.

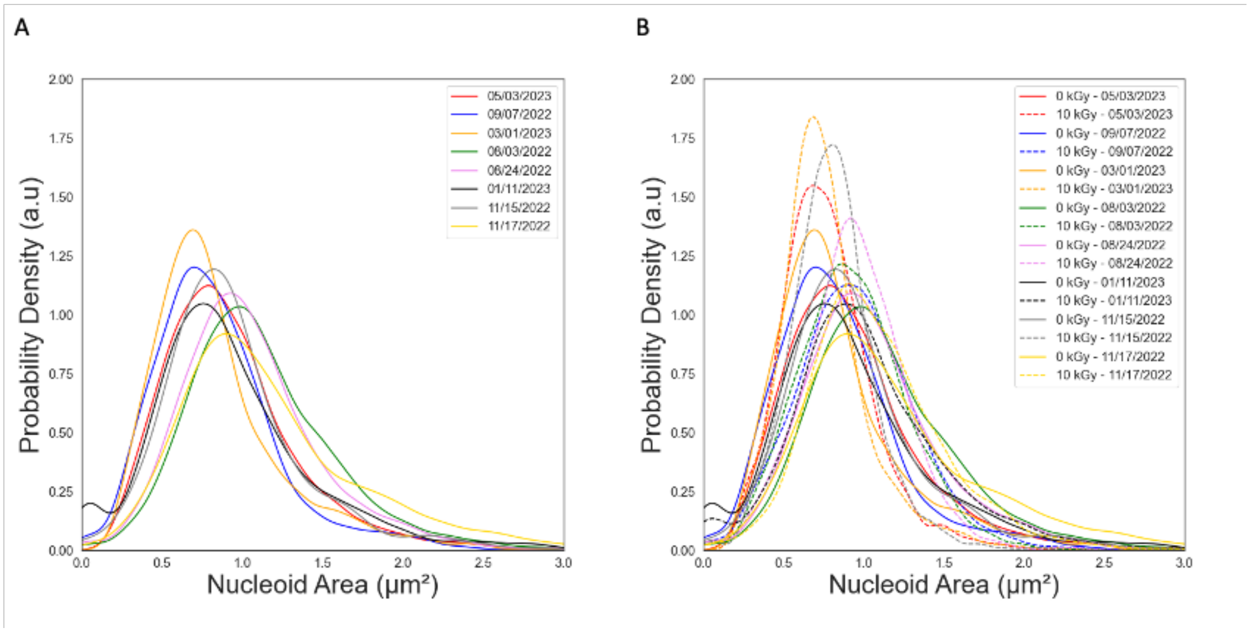


**Supplementary Figure S3.** Probability distributions of nucleoid areas for eight experiments each composed of 3 biological replicates, measured on the dates indicated in the legend. **(A-B)** For each biological replicate, the probability distribution of nucleoid areas for the non-irradiated (control) sample is shown with a solid line, and **(B)** the probability distribution of nucleoid areas for the sample irradiated at 10 kGy is shown with a dashed line of the same color.

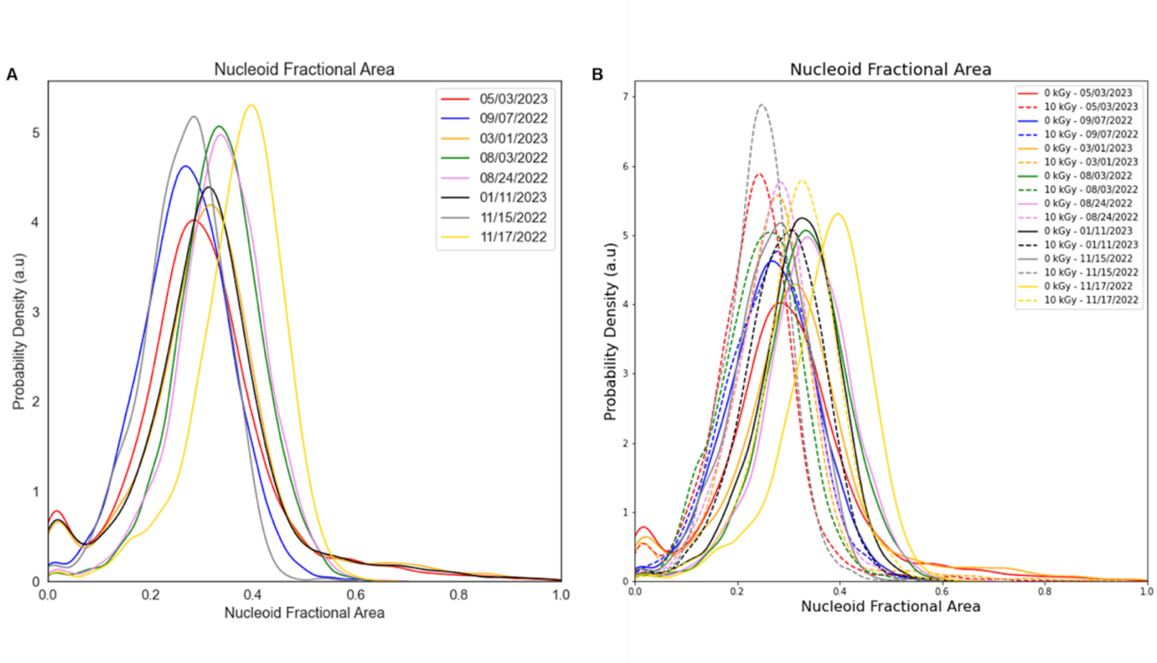


**Supplementary Figure S4.** Probability distributions of nucleoid fractional areas for eight experiments each composed of 3 biological replicates, measured on the dates indicated in the legend. **(A-B)** For each biological replicate, the probability distribution of nucleoid fractional areas for the non-irradiated (control) sample is shown with a solid line, and **(B)** the probability distribution of nucleoid fractional areas for the sample irradiated at 10 kGy is shown with a dashed line of the same color.

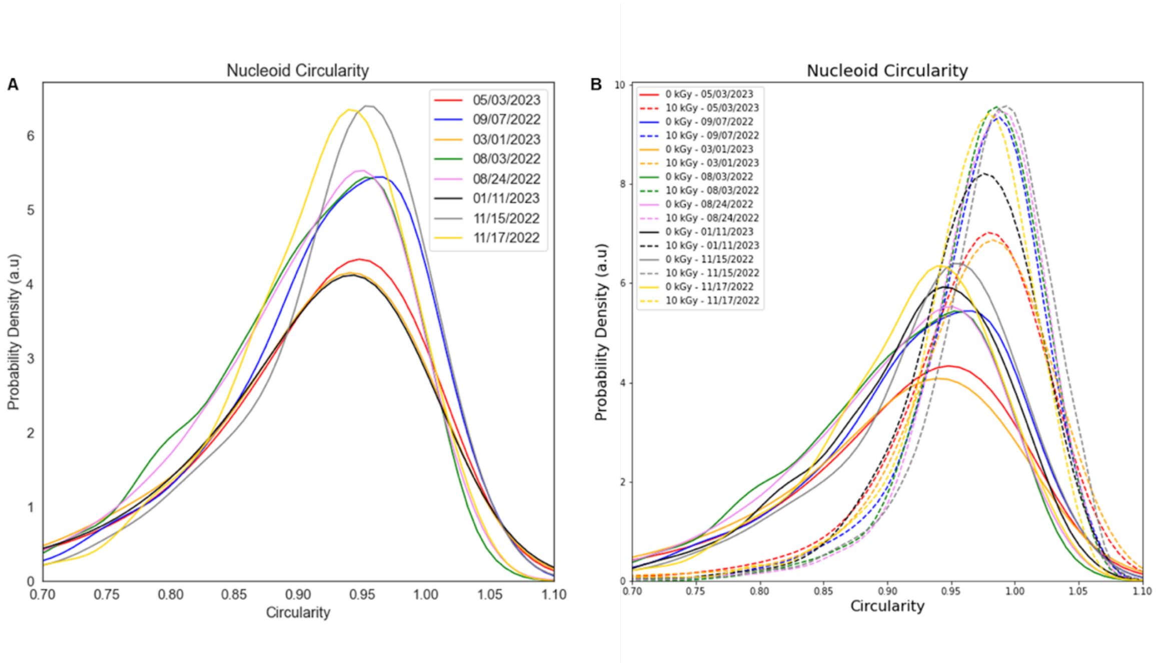


**Supplementary Figure S5.** Probability distributions of nucleoid circularities for eight experiments each composed of 3 biological replicates, measured on the dates indicated in the legend. **(A-B)** For each biological replicate, the probability distribution of nucleoid circularity for the non-irradiated (control) sample is shown with a solid line, and **(B)** the probability distribution of nucleoid circularity for the sample irradiated at 10 kGy is shown with a dashed line of the same color.

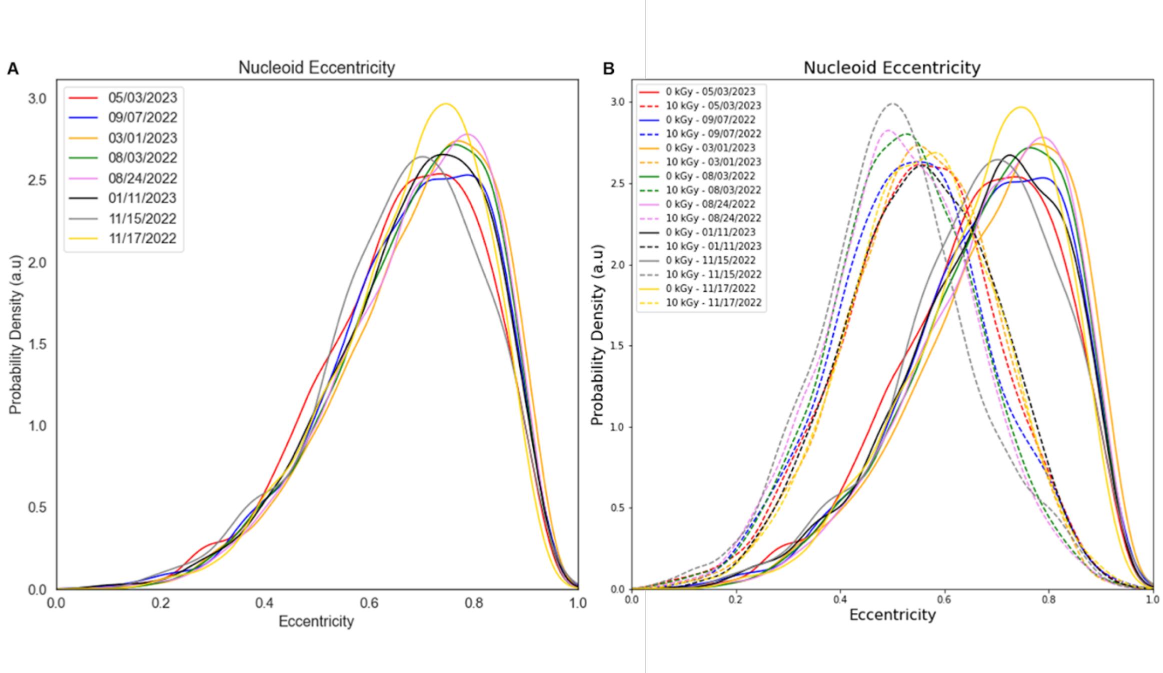


**Supplementary Figure S6.** Probability distributions of nucleoid eccentricity for eight experiments each composed of 3 biological replicates, measured on the dates indicated in the legend. **(A-B)** For each biological replicate, the probability distribution of nucleoid eccentricity for the non-irradiated (control) sample is shown with a solid line, and **(B)** the probability distribution of nucleoid eccentricity for the sample irradiated at 10 kGy is shown with a dashed line of the same color.

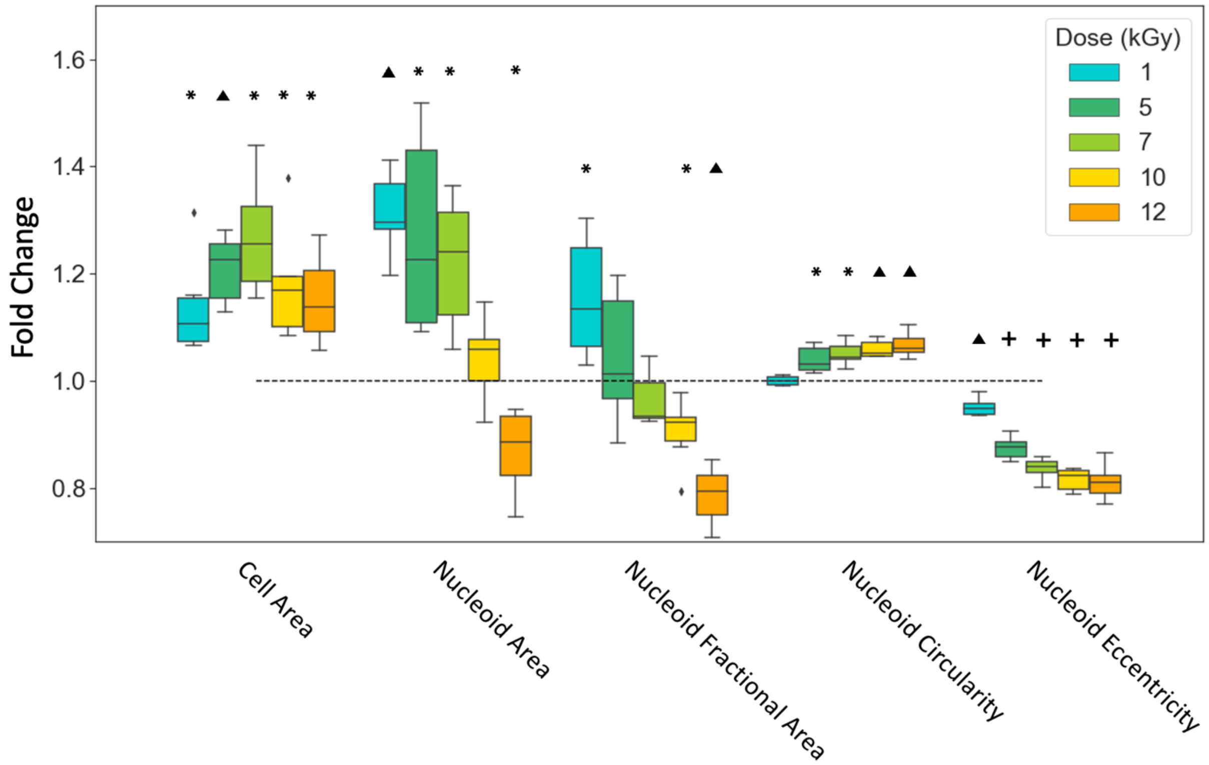


**Supplementary Figure S7.** Changes in the geometric characteristics of *D. radiodurans* cells and nucleoids following IR. Fold changes of the mean values of geometric properties from the value measured at 0 kGy. Measured images were taken after 2 hours of recovery. Each box-and-whisker plot contains data from 6 biological replicates over 2 experimental days. P-values are calculated using a student T-test with the null hypothesis that the fold change is equal to 1 (indicated by the black dashed horizontal line). * = $p < 0.05$, ▲ = $p < 0.001$, ✚ = $p < 0.0001$.

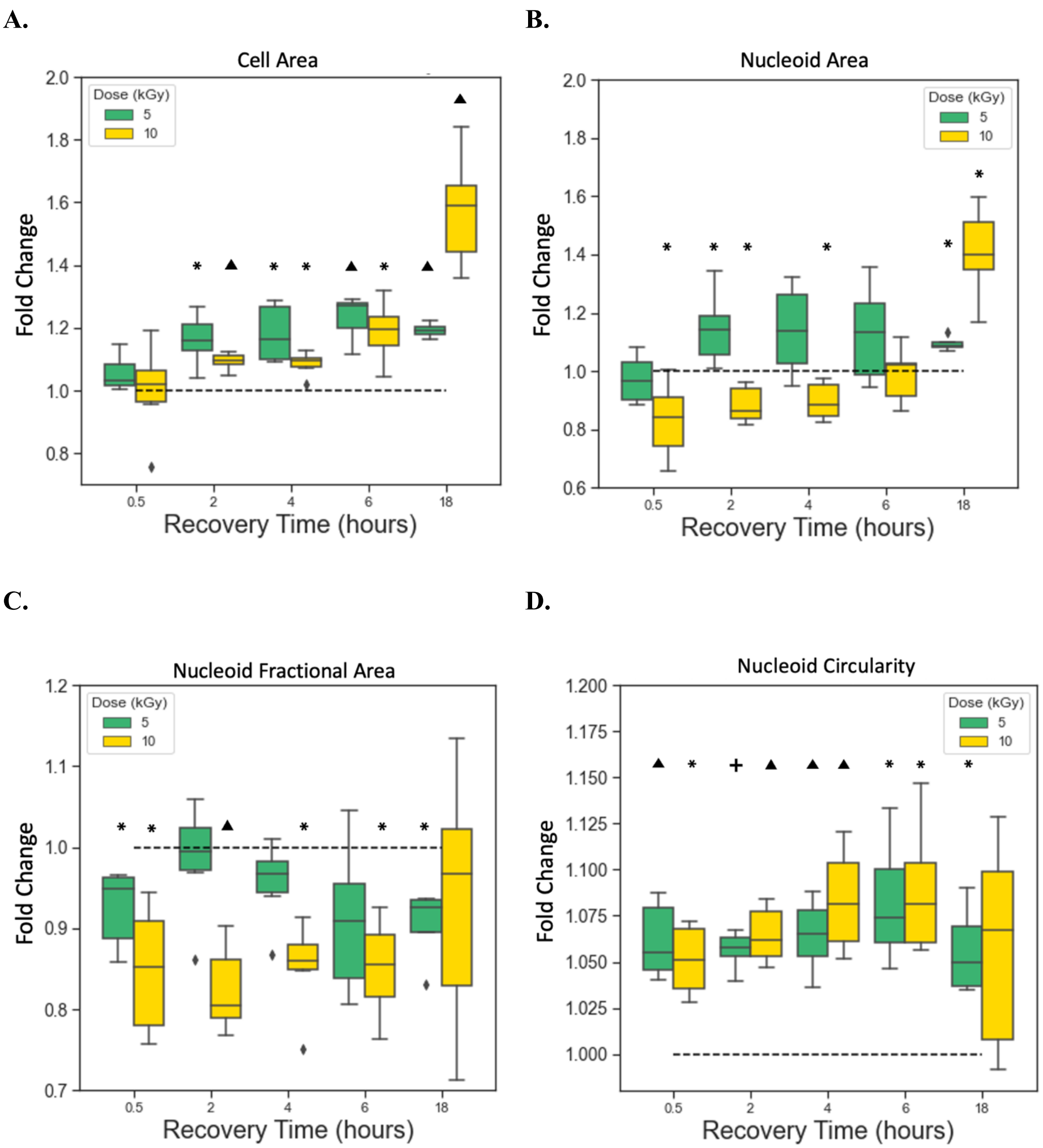


**Supplementary Figure S8.** Fold changes of the mean values of geometric characteristics from the value measured at 0 kGy for the same biological replicate, at the corresponding recovery time point. Each box-and-whisker plot contains data from 6 biological replicates over 2 days of experiments. Shown are **(A)** cell area, **(B)** nucleoid area, **(C)** nucleoid fractional area, and **(D)** nucleoid circularity. **(A-D)** P-values are calculated using a student T-test with the null hypothesis that the fold change is equal to 1 (indicated by the black dashed horizontal line). * = $p < 0.05$, ▲ = $p < 0.001$, ✚ = $p < 0.0001$.

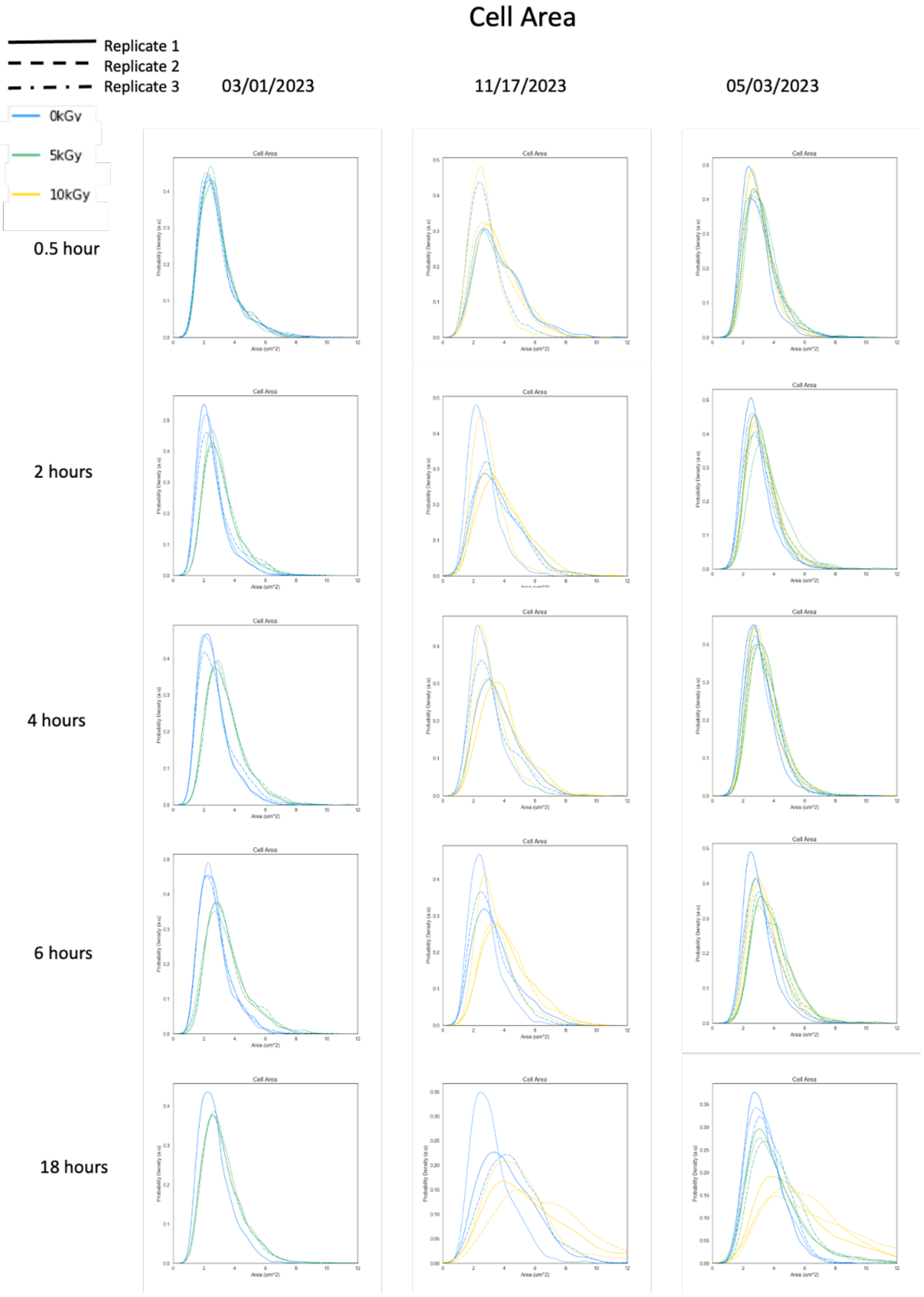
Cell Area
Replicate 1
Replicate 2
Replicate 3
0kGy
5kGy
10kGy
03/01/2023
11/17/2023
05/03/2023
0.5 hour
2 hours
4 hours
6 hours
18 hours
Cell Area
Area (um^2)
Probability Density (a.u)

**Supplementary Figure S9.** Probability distributions of cell area measured for the date indicated at the top of each column. Each date has three biological replicates, distinguished by a solid line, a dashed line, and a dashed-dot line. Radiation dose is indicated by the color of the line, as indicated in the legend. The recovery time at which each set of measurements was taken is indicated by the times indicated to the left of each row. In total, $N = 6$ for irradiated samples, $N = 9$ for non-irradiated controls.

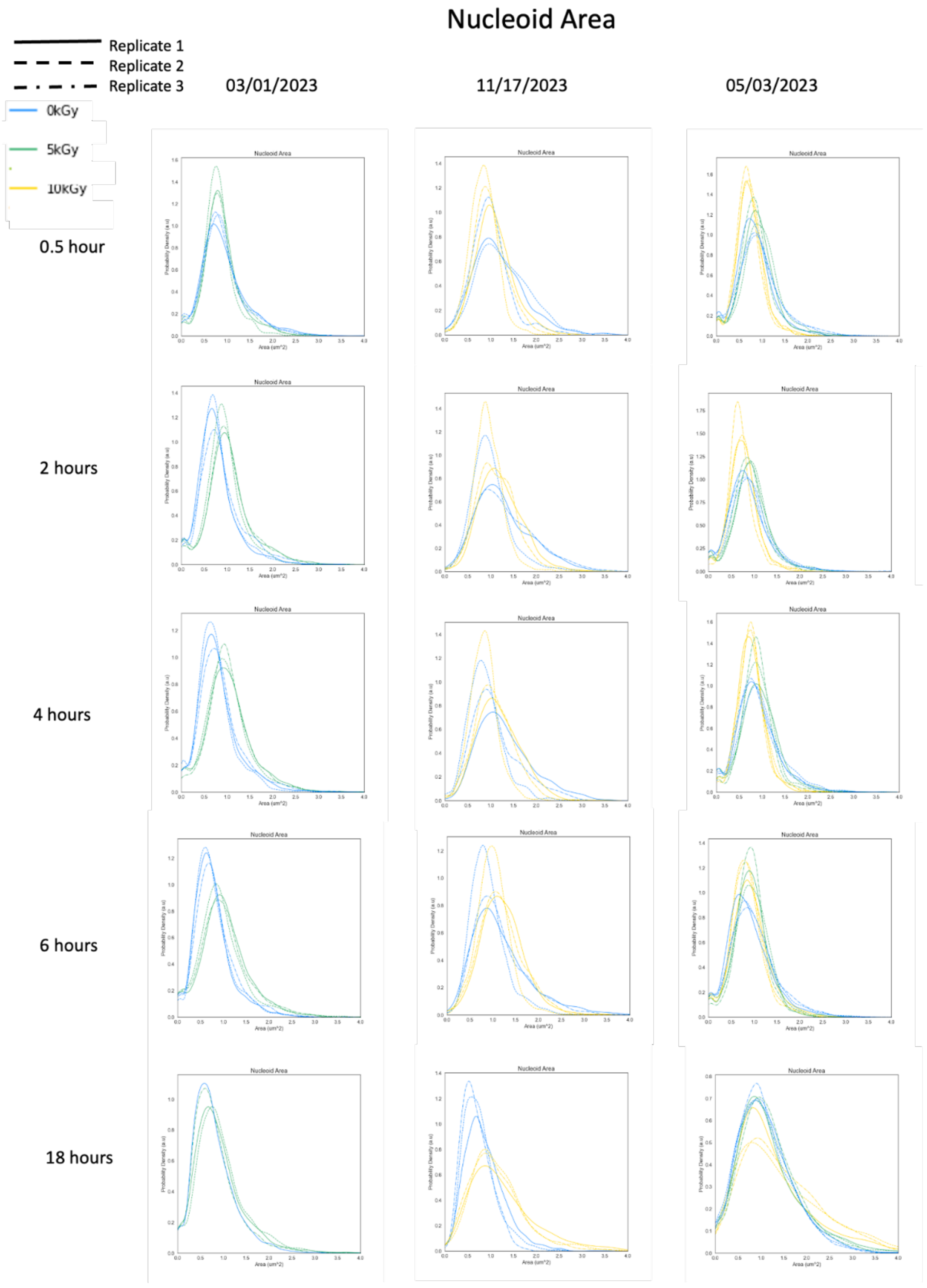
Nucleoid Area
Replicate 1
Replicate 2
Replicate 3
0kGy
5kGy
10kGy
03/01/2023
11/17/2023
05/03/2023
0.5 hour
2 hours
4 hours
6 hours
18 hours
Probability Density (a.u)
Area (um^2)

**Supplementary Figure S10.** Probability distributions of nucleoid area measured for the date indicated at the top of each column. Each date has three biological replicates, distinguished by a solid line, a dashed line, and a dashed-dot line. Radiation dose is indicated by the color of the line, as indicated in the legend. The recovery time at which each set of measurements was taken is indicated by the times indicated to the left of each row. In total, $N = 6$ for irradiated samples, $N = 9$ for non-irradiated controls.

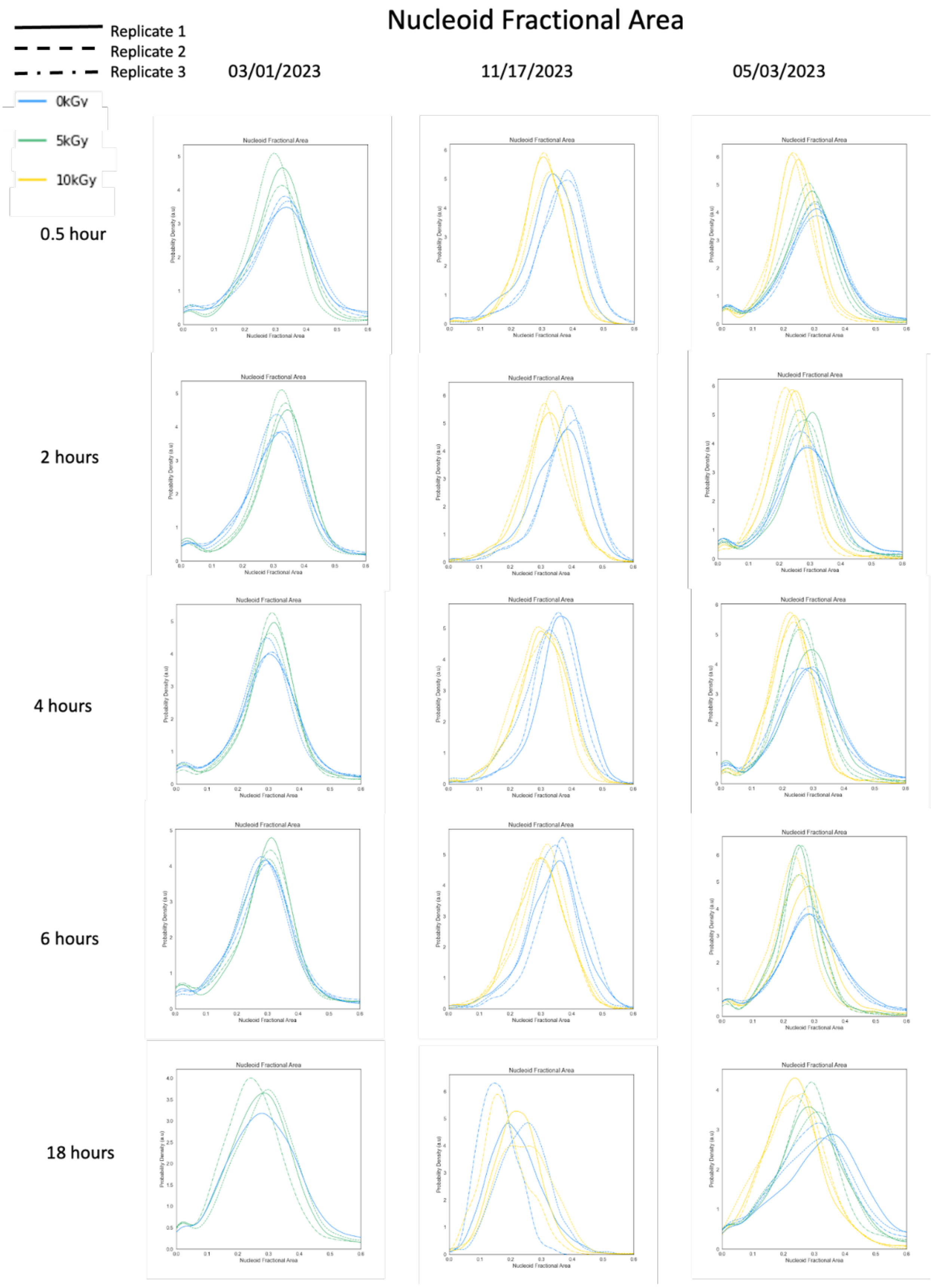

Nucleoid Fractional Area
Replicate 1
Replicate 2
Replicate 3
0kGy
5kGy
10kGy
03/01/2023
11/17/2023
05/03/2023
0.5 hour
2 hours
4 hours
6 hours
18 hours

**Supplementary Figure S11.** Probability distributions of nucleoid fractional area measured for the date indicated at the top of each column. Each date has three biological replicates, distinguished by a solid line, a dashed line, and a dashed-dot line. Radiation dose is indicated by the color of the line, as indicated in the legend. The recovery time at which each set of measurements was taken is indicated by the times indicated to the left of each row. In total, $N = 6$ for irradiated samples, $N = 9$ for non-irradiated controls.

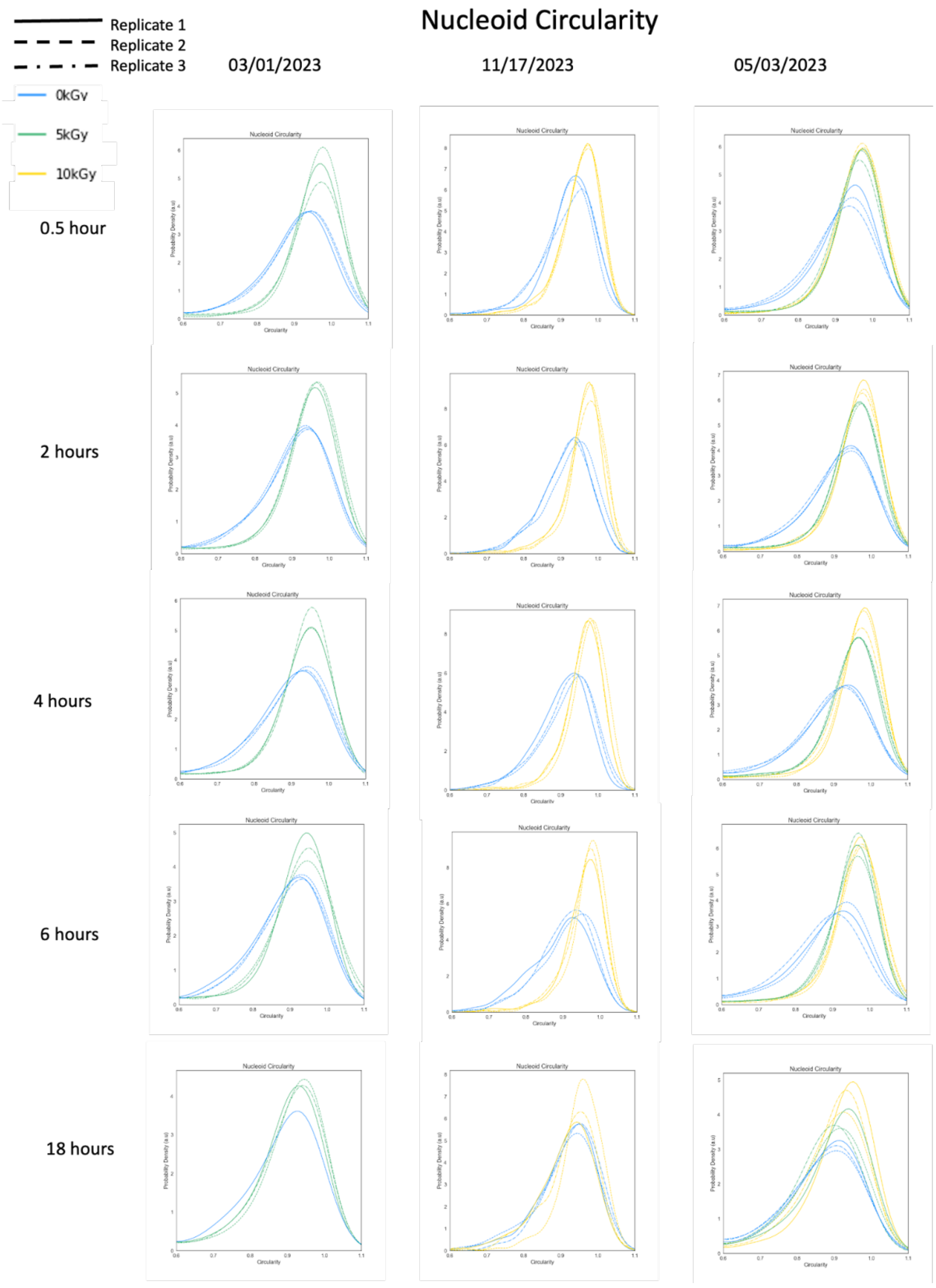

Nucleoid Circularity
Replicate 1
Replicate 2
Replicate 3
03/01/2023
11/17/2023
05/03/2023
0kGy
5kGy
10kGy
0.5 hour
2 hours
4 hours
6 hours
18 hours
Nucleoid Circularity
Probability Density (a.u)
Circularity

**Supplementary Figure S12.** Probability distributions of nucleoid circularity measured for the date indicated at the top of each column. Each date has three biological replicates, distinguished by a solid line, a dashed line, and a dashed-dot line. Radiation dose is indicated by the color of the line, as indicated in the legend. The recovery time at which each set of measurements was taken is indicated by the times indicated to the left of each row. In total, $N = 6$ for irradiated samples, $N = 9$ for non-irradiated controls.

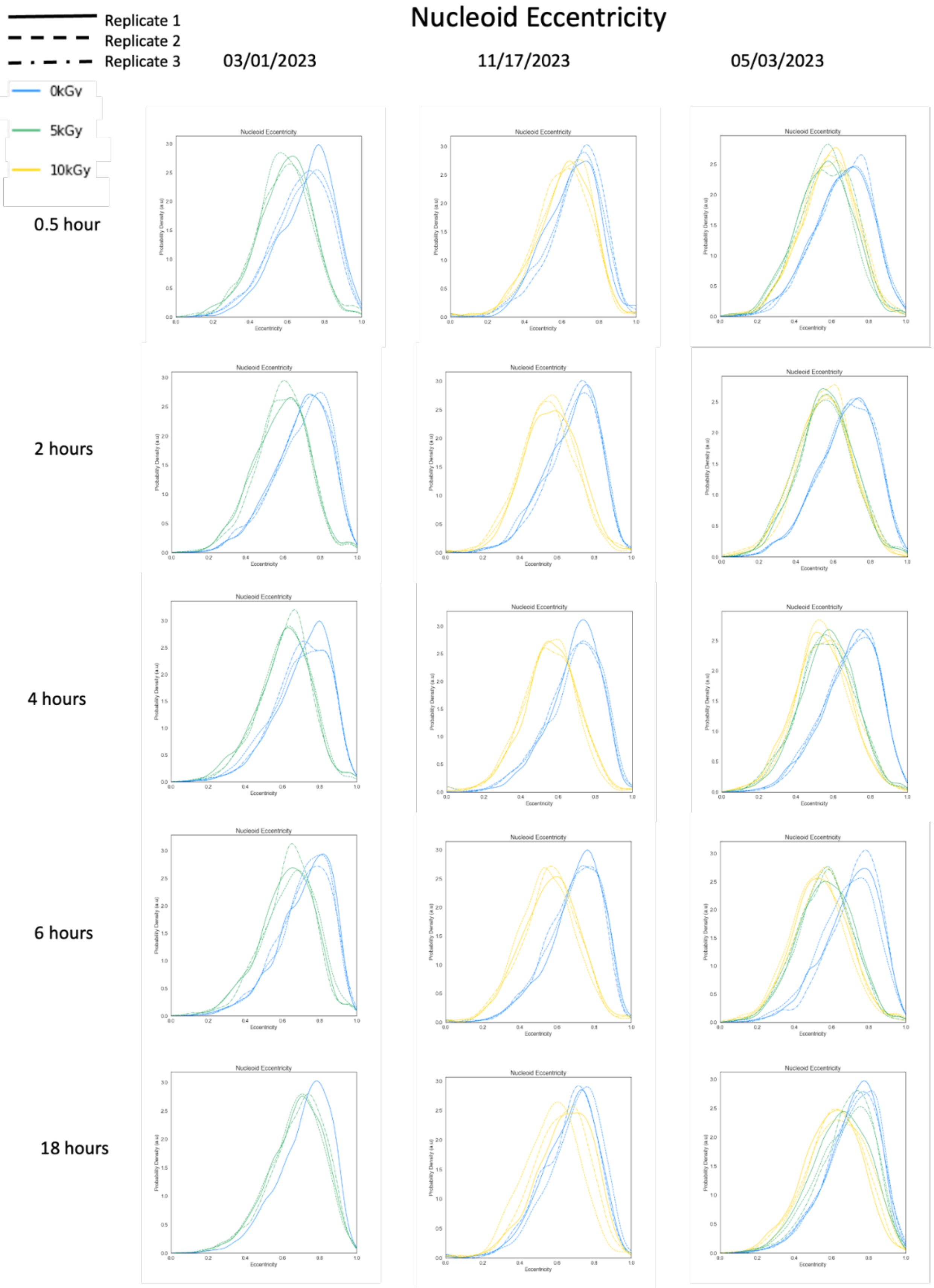
Nucleoid Eccentricity
Replicate 1
Replicate 2
Replicate 3
0kGy
5kGy
10kGy
03/01/2023
11/17/2023
05/03/2023
0.5 hour
2 hours
4 hours
6 hours
18 hours

**Supplementary Figure S13.** Probability distributions of nucleoid eccentricity measured for the date indicated at the top of each column. Each date has three biological replicates, distinguished by a solid line, a dashed line, and a dashed-dot line. Radiation dose is indicated by the color of the line, as indicated in the legend. The recovery time at which each set of measurements was taken is indicated by the times indicated to the left of each row. In total, $N = 6$ for irradiated samples, $N = 9$ for non-irradiated controls.

**A.** **B.**

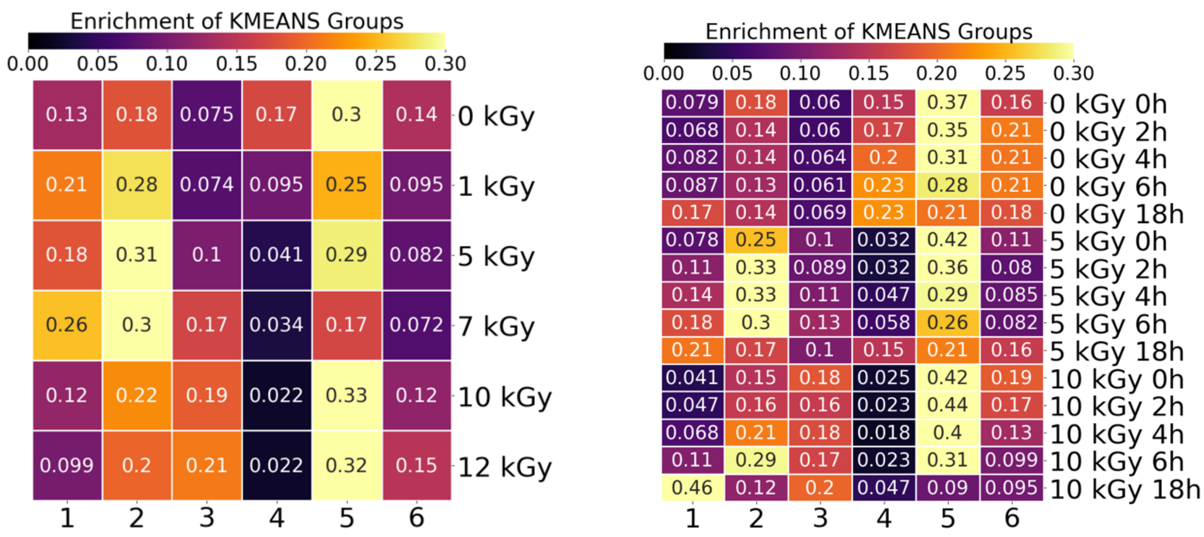


**Supplementary Figure S14.** Enrichment of the k-means clusters for the wild-type *D. radiodurans* strain. **A.** k-means values of the *D. radiodurans* wild-type strain at 0, 1, 5, 7, 10, and 12 kGy respectively. **B.** k-means values of the *D. radiodurans* wild-type strain over an 18-hour recovery period at the 0, 5, and 10 kGy acute ionizing radiation doses respectively.

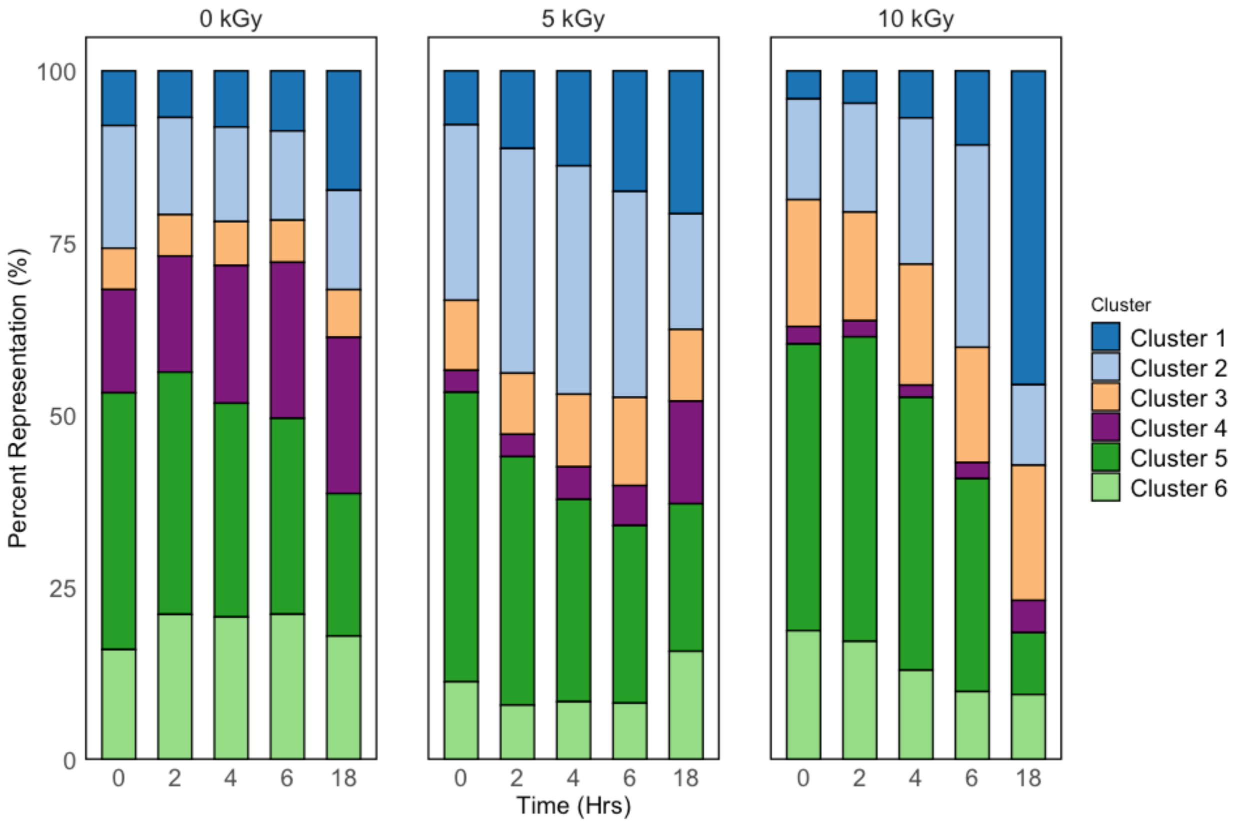


**Supplementary Figure S15.** Shifts in morphological cluster representation with acute IR dose treatments for the wild-type *D. radiodurans* strain over time. Cluster representation is displayed as a percentage of the total population of cells contained within each cluster defined in Figure 4A.

**A.**

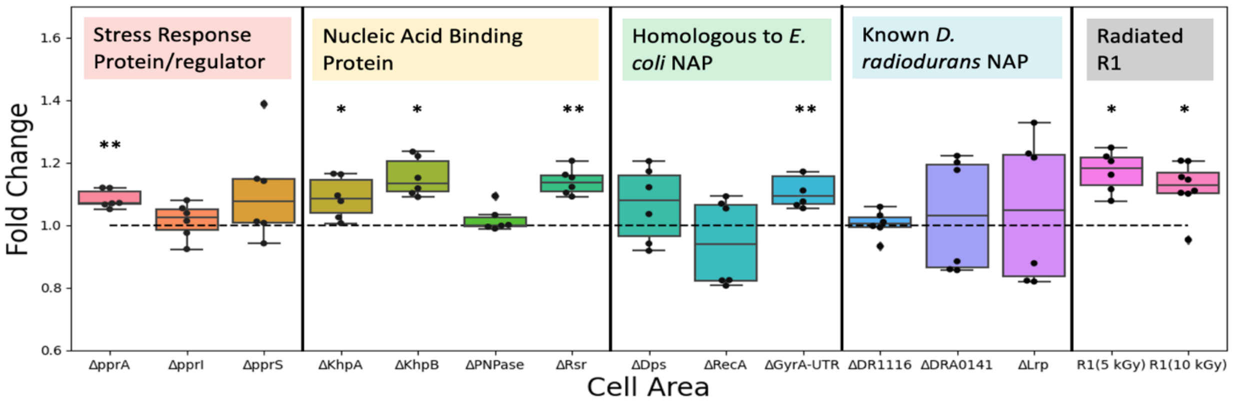

Stress Response Protein/regulator
Nucleic Acid Binding Protein
Homologous to *E. coli* NAP
Known *D. radiodurans* NAP
Radiated R1
Fold Change
1.6
1.4
1.2
1.0
0.8
0.6
**
*
*
**
**
*
*
ΔpprA
ΔpprI
ΔpprS
ΔKhpA
ΔKhpB
ΔPNPase
ΔRsr
ΔDps
ΔRecA
ΔGyrA-UTR
ΔDR1116
ΔDRA0141
ΔLrp
R1(5 kGy)
R1(10 kGy)
Cell Area


**B.**

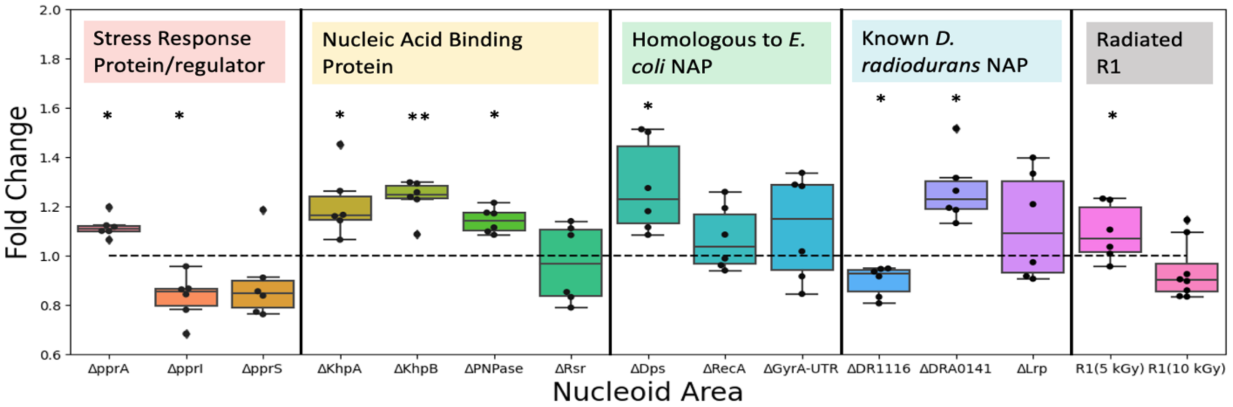

Stress Response Protein/regulator
Nucleic Acid Binding Protein
Homologous to *E. coli* NAP
Known *D. radiodurans* NAP
Radiated R1
Fold Change
2.0
1.8
1.6
1.4
1.2
1.0
0.8
0.6
*
*
*
**
*
*
*
*
*
ΔpprA
ΔpprI
ΔpprS
ΔKhpA
ΔKhpB
ΔPNPase
ΔRsr
ΔDps
ΔRecA
ΔGyrA-UTR
ΔDR1116
ΔDRA0141
ΔLrp
R1(5 kGy)
R1(10 kGy)
Nucleoid Area


**C.**

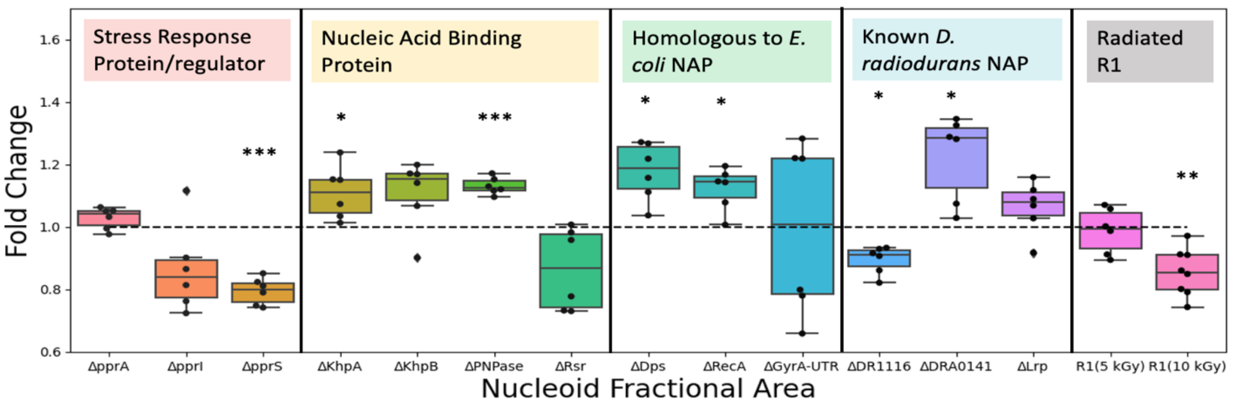

Stress Response Protein/regulator
Nucleic Acid Binding Protein
Homologous to *E. coli* NAP
Known *D. radiodurans* NAP
Radiated R1
Fold Change
1.6
1.4
1.2
1.0
0.8
0.6
***
*
***
*
*
*
*
**
ΔpprA
ΔpprI
ΔpprS
ΔKhpA
ΔKhpB
ΔPNPase
ΔRsr
ΔDps
ΔRecA
ΔGyrA-UTR
ΔDR1116
ΔDRA0141
ΔLrp
R1(5 kGy)
R1(10 kGy)
Nucleoid Fractional Area

**D.**

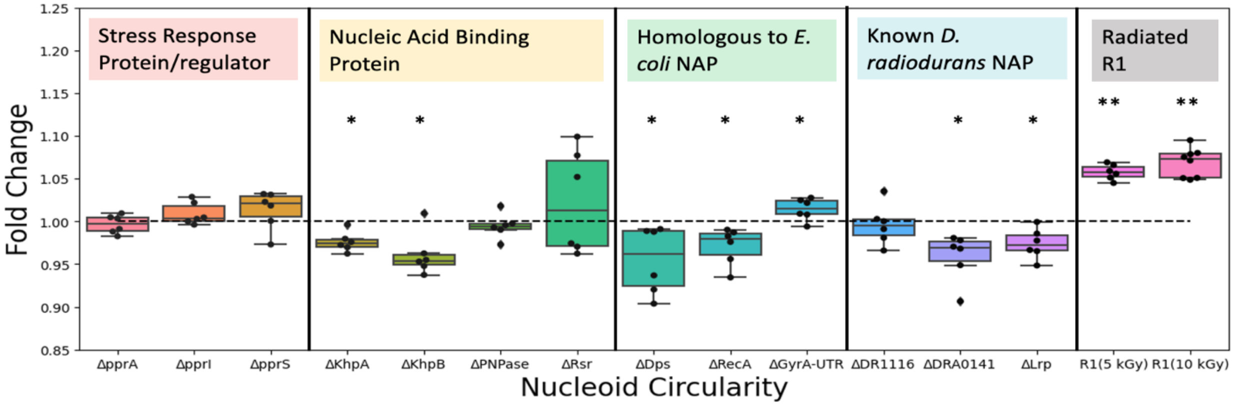


**E.**

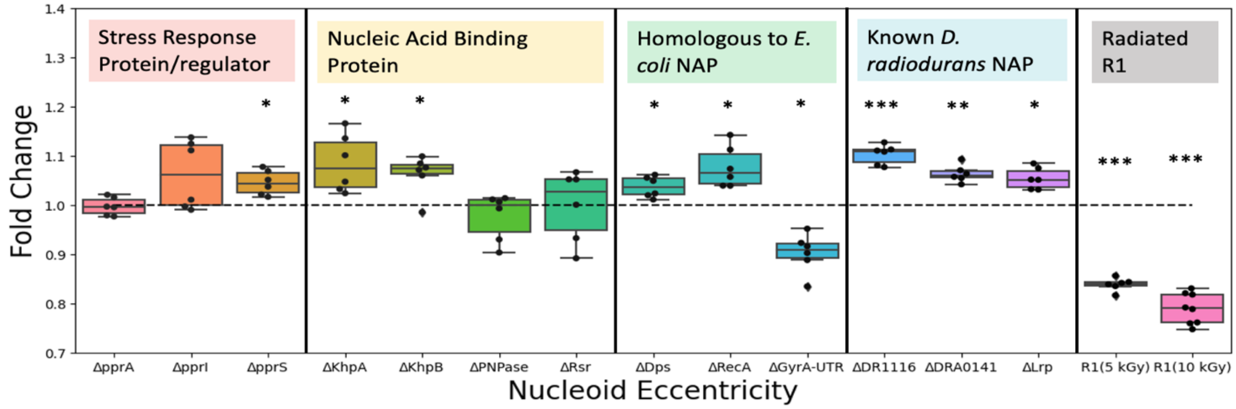


**Supplementary Figure S16.** Effects of genomic protein knockouts on geometric properties, measured as fold changes of the mean geometric value for the mutant to the corresponding mean geometric value for wild-type R1 *D. radiodurans*. The horizontal dashed lines correspond to fold changes of one, meaning no difference between the mutant and the wild-type for that measured property. Shown are **(A)** cell area, **(B)** nucleoid area, **(C)** nucleoid fractional area, **(D)** nucleoid circularity, and **(E)** nucleoid eccentricity. For each genomic mutant, six biological replicates were used (N = 6). Statistical significance in the fold changes (with a null hypothesis of a fold change of one) is indicated by * = $p < 0.05$, ** = $p < 0.001$, *** = $p < 0.0001$ and recorded in Supplementary Table S9.

**A.** **B.**

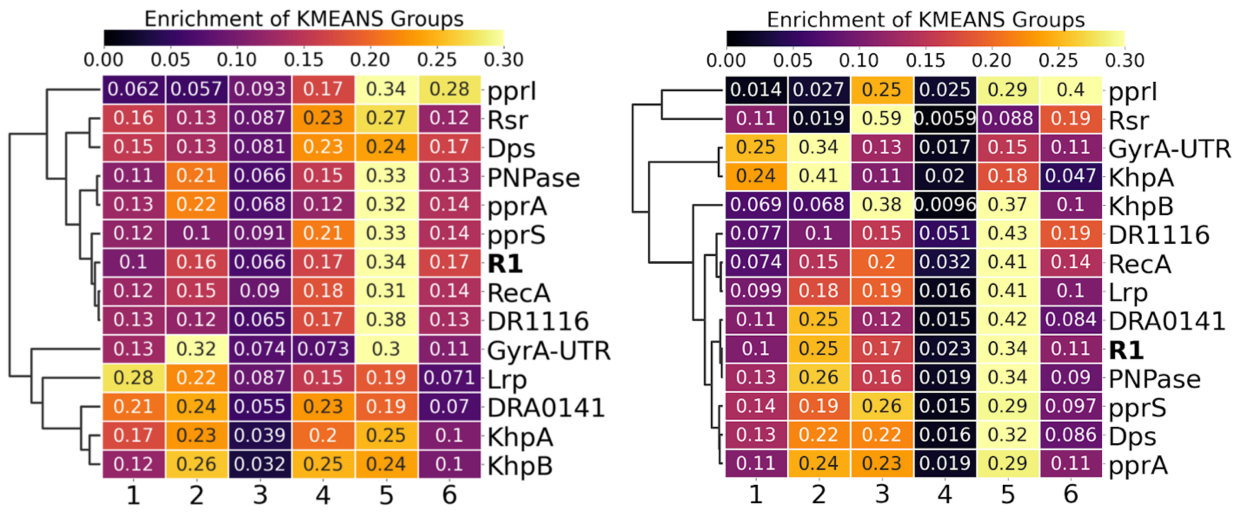


**Supplementary Figure S17.** Enrichment of the k-means clusters for *D. radiodurans* strains. **A.** k-means values of the *D. radiodurans* wild-type strain (R1) and genomic mutant strains at 0 kGy of ionizing radiation. **B.** k-means values of the *D. radiodurans* wild-type strain (R1) and genomic mutant strains at 10 kGy of ionizing radiation.

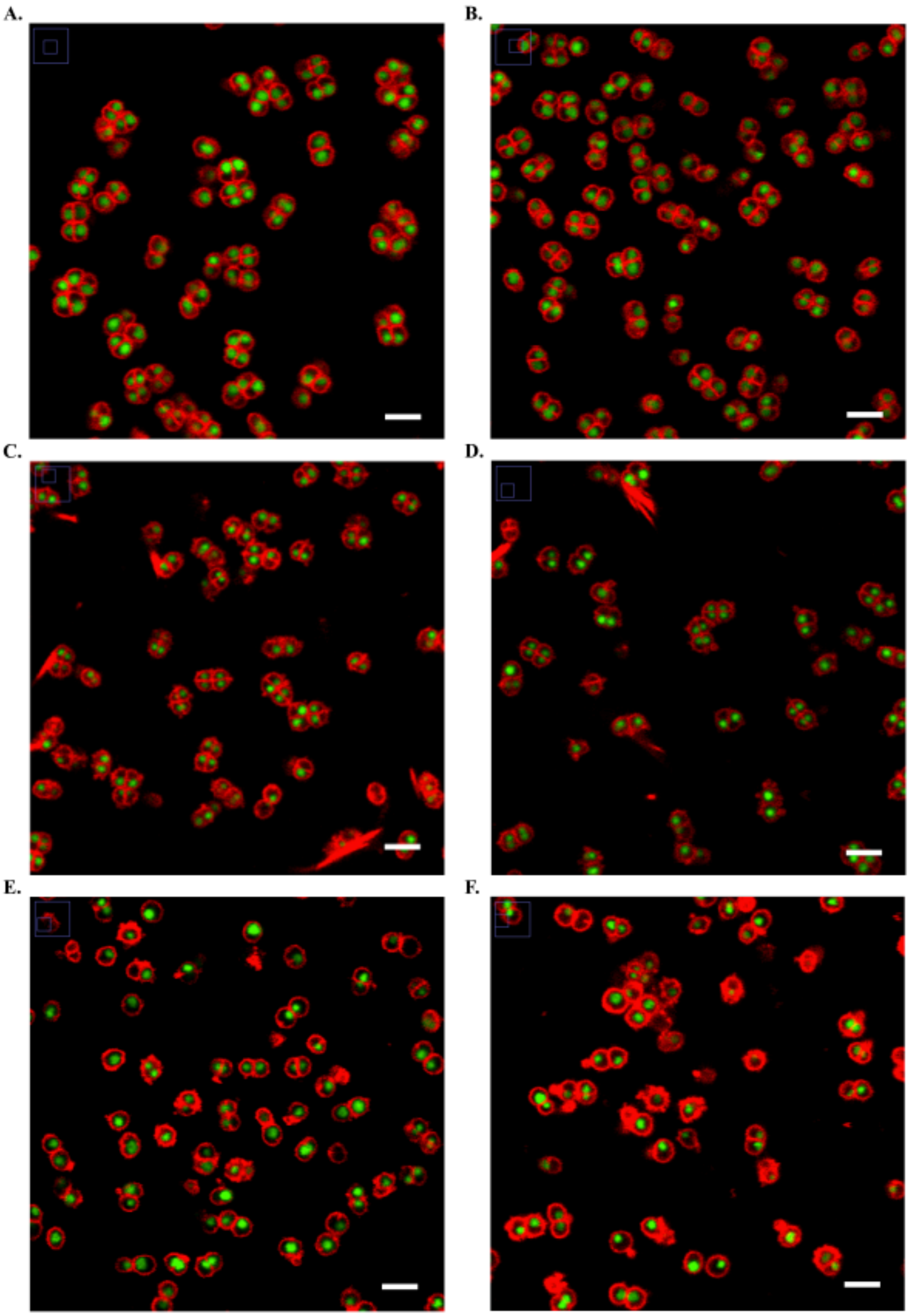


**Supplementary Figure S18.** *ΔGyrA-UTR* blebbing. Strains were exposed to 10 kGy acute IR and were imaged 2-hour post recovery. Biological replicates shown are from two experiment days. Cells are stained as follows – Red: Nile Red stain (membrane), Green: Syto 9 stain (nucleoid). **A.** R1 control; day 1. **B.** R1 control; day 2. **C.** *ΔGyrA-UTR* strain; day 1; biological replica A. **D.** *ΔGyrA-UTR* strain; day 1; biological replica B. **E.** *ΔGyrA-UTR* strain; day 2; biological replica A. **F.** *ΔGyrA-UTR* strain; day 2; biological replica B. Scale bar: 5 μm.

**A.** **B.** **C.**

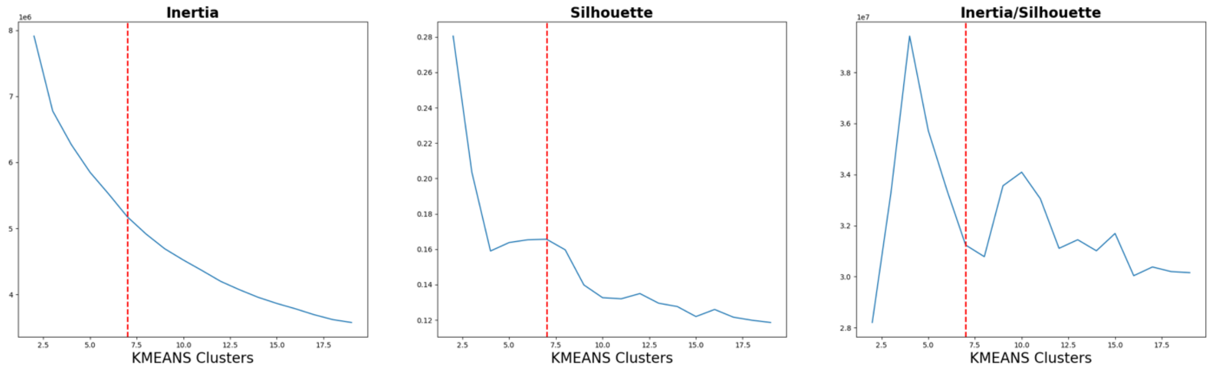


**Supplementary Figure S19.** Algorithmic convergence on six optimal clusters for k-means analysis based on the “elbow method”, or a plateau of inertia/silhouette values. (A) Inertia values as a function of the number of clusters. (B) Silhouette values as a function of the number of clusters. (C) Inertia/silhouette value begins to plateau at approximately 6 clusters, given a constraint of number of clusters being greater than two.
